# Structure-based prediction of HDAC6 substrates validated by enzymatic assay reveals determinants of promiscuity and detects new potential substrates

**DOI:** 10.1101/2021.02.21.431878

**Authors:** Julia K. Varga, Kelsey Diffley, Katherine R. Welker Leng, Carol A. Fierke, Ora Schueler-Furman

## Abstract

Histone deacetylases play important biological roles well beyond the deacetylation of histone tails. In particular, HDAC6 is involved in multiple cellular processes such as apoptosis, cytoskeleton reorganization, and protein folding, affecting substrates such as α-tubulin, Hsp90 and cortactin proteins. We have applied a biochemical enzymatic assay to measure the activity of HDAC6 on a set of candidate unlabeled peptides. These served for the calibration of a structure-based substrate prediction protocol, Rosetta FlexPepBind, previously used for the successful substrate prediction of HDAC8 and other enzymes. A proteome-wide screen of reported acetylation sites using our calibrated protocol together with the enzymatic assay provide new peptide substrates and avenues to novel potential functional regulatory roles of this promiscuous, multi-faceted enzyme. In particular, we propose novel regulatory roles of HDAC6 in tumorigenesis and cancer cell survival via the regulation of the EGFR/Akt pathway activation process. The calibration process and comparison of the results between HDAC6 and HDAC8 highlight structural differences that explain the established promiscuity of HDAC6.

## Introduction

In order for cells and organisms to survive and adapt to different conditions, complex, tightly-controlled, and context-dependent regulation is crucial. Much of this regulation is achieved by post-translational modifications (PTMs) that can change the behavior of a protein (as well as corresponding modifications of other macromolecules such as DNA and RNA). One of the major regulatory modifications is the acetylation and deacetylation of histones, which is a main component of the histone code dictating chromatin organization and transcriptional activity^1,2^, mediated by lysine acetylases and deacetylases (HATs and HDACs, respectively). HDACs catalyze the removal of an acetyl group from the post-translational modification of acetyl-lysine in proteins.

Lysine deacetylase 6 (HDAC6) is a class IIB Zn^2+^ deacetylase and is the only HDAC to contain two deacetylase domains of distinct specificities. The first domain specifically deacetylates acetylated C-terminal lysine residues, while the second shows a particularly broad substrate selectivity^3,4^. There is evidence that HDAC6 catalyzes deacetylation of several proteins involved in a variety of cellular processes. Among them, HDAC6-mediated deacetylation of α-tubulin regulates microtubule stability and cell motility^5^. Another characterized substrate, cortactin, binds to deacetylated actin filaments and participates in the fusion of lysosomes and autophagosomes^6^. The enzyme also plays a role in protein folding by regulating the activity of the Hsp90 chaperone protein via deacetylation^7,8^. In addition, HDAC6 is an important player in innate immunity, regulating the detection of pathogen genomic material via deacetylation of retinoic acid inducible gene-I protein^9–11^. While the broad specificity of HDAC6 has been reported, a full understanding of the selectivity determinants is still lacking, as is a proper understanding of the underlying structural basis that makes this particular HDAC more promiscuous than others, such as HDAC8^12^.

Beyond the few known substrates of HDAC6 mentioned above, substrate selectivity of human HDAC6 has been assessed at large scale in three key experimental studies^13,14,3^. Riester *et al*. used arrays of trimer peptides conjugated to 7-amino-4-methylcoumarin (AMC) and measured enzymatic activity by change in fluorescence^13^ (referred to as dataset D-3MER in this study). Schölz *et al*. applied specific inhibitors against several HDACs in cell lines and quantified the change in acetylated lysine sites by SILAC-MS (dataset D-SILAC)^14^. In both studies, the experiments were run with at least 5 different HDACs, and both reached the conclusion that HDAC6 was, by far, the most promiscuous among the examined HDACs. Finally, Kutil *et al*. assessed HDAC6 deacetylation of 13-mer peptides synthesized on an array, measuring activity with a mixture of anti-acetyl lysine antibodies (dataset D-13MER) and verified 20 substrate hits by gas chromatography-mass spectrometry (GC-MS; dataset D-GCMS)^3^. In the latter study, the predicted substrates were also compared with hits from other studies, finding minimal overlap^3^ (see **Table S1** for a summary of all studies of HDAC6 substrates).

Many of the enzymes that add or remove PTMs act on short linear motifs (SLIMs) that are often exposed. Therefore, their substrate selectivity may be approximated by short peptides that cover the region^15^. Different types of prediction methods have been developed for finding putative substrates. Many sequence-based predictions find modification sites based on position-specific scoring matrices (PSSMs) or regular expressions^16^ that are derived from a large set of substrates, often selected by high-throughput approaches, such as those described above. However, these do not account for possible interdependencies between amino acids at different positions in the substrate, nor do they consider secondary structure that might be important for recognition. Machine Learning-based approaches can be used for these aims (e.g. Hidden Markov Models^17^ and naive Bayes^18^), but such approaches depend on considerable amounts of data^19–21^. Moreover, enzyme substrate patterns may not always adequately be depicted by a sequence-based description, like in the case of O-glycosylation^22^ or HIV-1 protease substrates^23^.

Structure-based methods can complement sequence-based methods, particularly in cases of non-canonical motifs^22,24^, as we have previously shown for PTM enzymes using the Rosetta FlexPepBind protocol^12,25,26^. This approach assumes that the ability of the substrate local peptide sequence to bind in a catalysis-competent conformation is a main determinant of enzyme selectivity, and thus the binding energy of such enzyme-substrate complex structures can be taken as a proxy for substrate activity. The accuracy of the calibrated protocol can be estimated by applying it to an independent test set. New substrates can then be identified by applying the calibrated protocol to candidate peptides with unknown activity.

In this study we utilized an accurate biochemical assay that measures acetate production following deacetylation^27^ to quantify the catalytic activity of HDAC6 for specific peptides and establish a gold standard set of peptide substrates. Based on these activities, we calibrated FlexPepBind to evaluate activity of potential substrates, as we have done in the past for other HDACs and PTM enzymes (e.g. HDAC8 and FTase^12,26^). Calibration revealed structural differences between HDAC6 and HDAC8 that form the basis of the considerable difference in selectivity of these two deacetylases. Application of this method to screen the acetylome identified novel potential regulatory mechanisms based on HDAC6-dependent regulation. In the end, the combination of our structure-based approach that is based on accurate, *in vitro* biochemical measures of substrate activity, with the previously reported large-scale approaches leads to a better understanding of HDAC6 substrate selectivity and biological function.

## Results

### Accurate measure of HDAC6 substrate activity using a biochemical enzymatic assay

We used an enzyme-coupled acetate detection assay, or simply the ‘acetate assay’ (see Methods), developed and applied previously^12,27^, to measure the catalytic efficiency (*k*_cat_/*K*_M_) of HDAC6. We measured catalysis of deacetylation of acetylated peptides with sequences taken from known substrate proteins, as well as a set of selected peptides with reported acetylation sites that were available to us from previous studies (**Table 1**). We used the second deacetylase domain (DD2) of HDAC6 for these experiments (from *Danio rerio*, which is more stable and has been shown to be a valid substitute for human HDAC6^4^). To ensure an accurate determination of *k*_cat_/*K*_M_ we measured HDAC6-catalyzed deacetylation at a minimum of four peptide concentrations, with at least two concentrations below *K*_M_ (**Figure 1**, squares). A total of 26 peptides which met these criteria formed our training set (D-TRAINING, **Table 1**). Additionally, we included 16 peptides where the value of *k*_cat_ was measured accurately but the *K*_M_ value was lower than the limit of detection for the acetate assay (∼10-20 μM substrate, circles in **Figure 1**) allowing only the determination of a lower limit for the value of *k*_cat_/*K*_M_ (set D-CAPPED, see below **Table 1**). The measured *k*_cat_/*K*_M_ values span the range of three orders of magnitude.

**Table 1.**
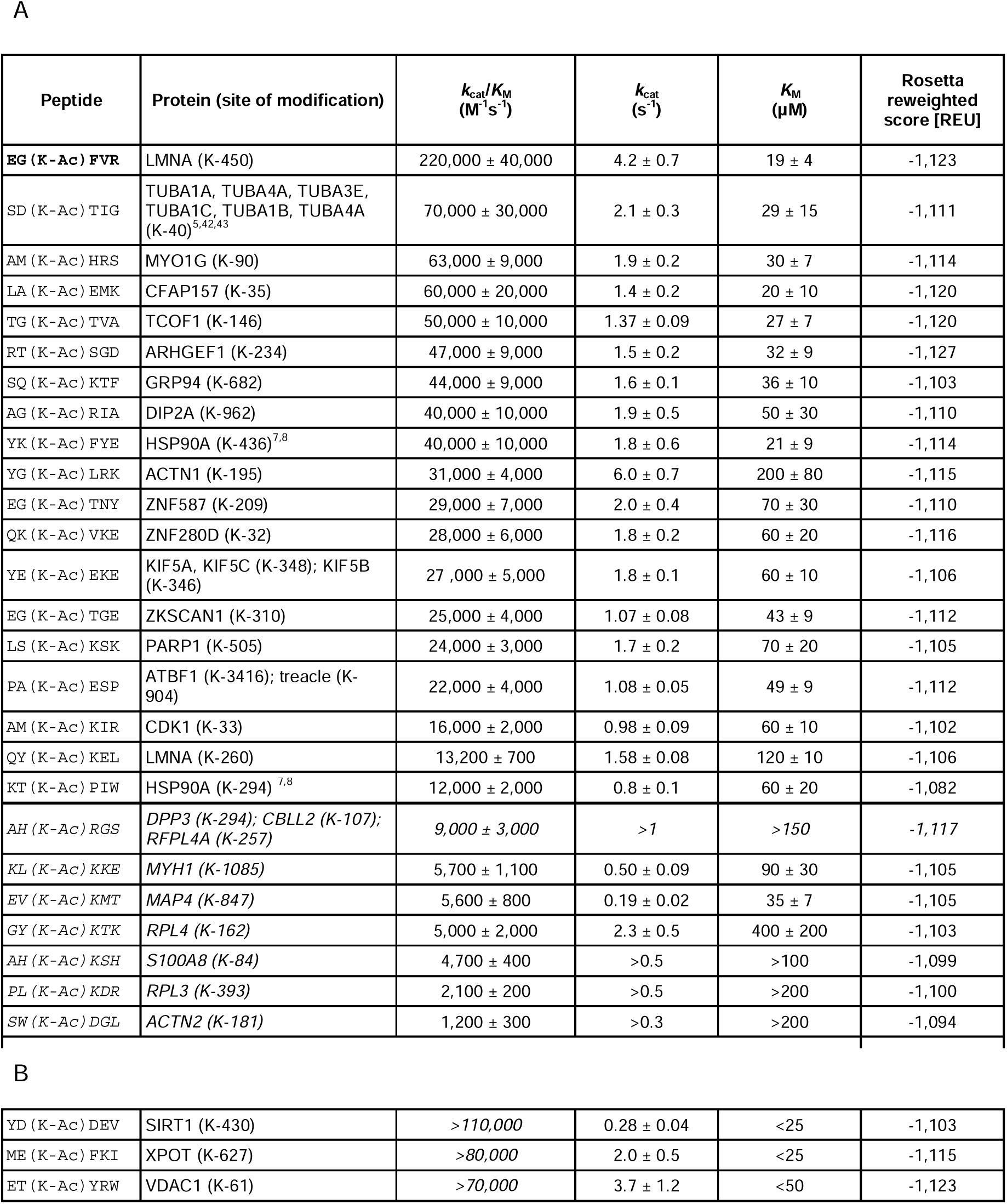
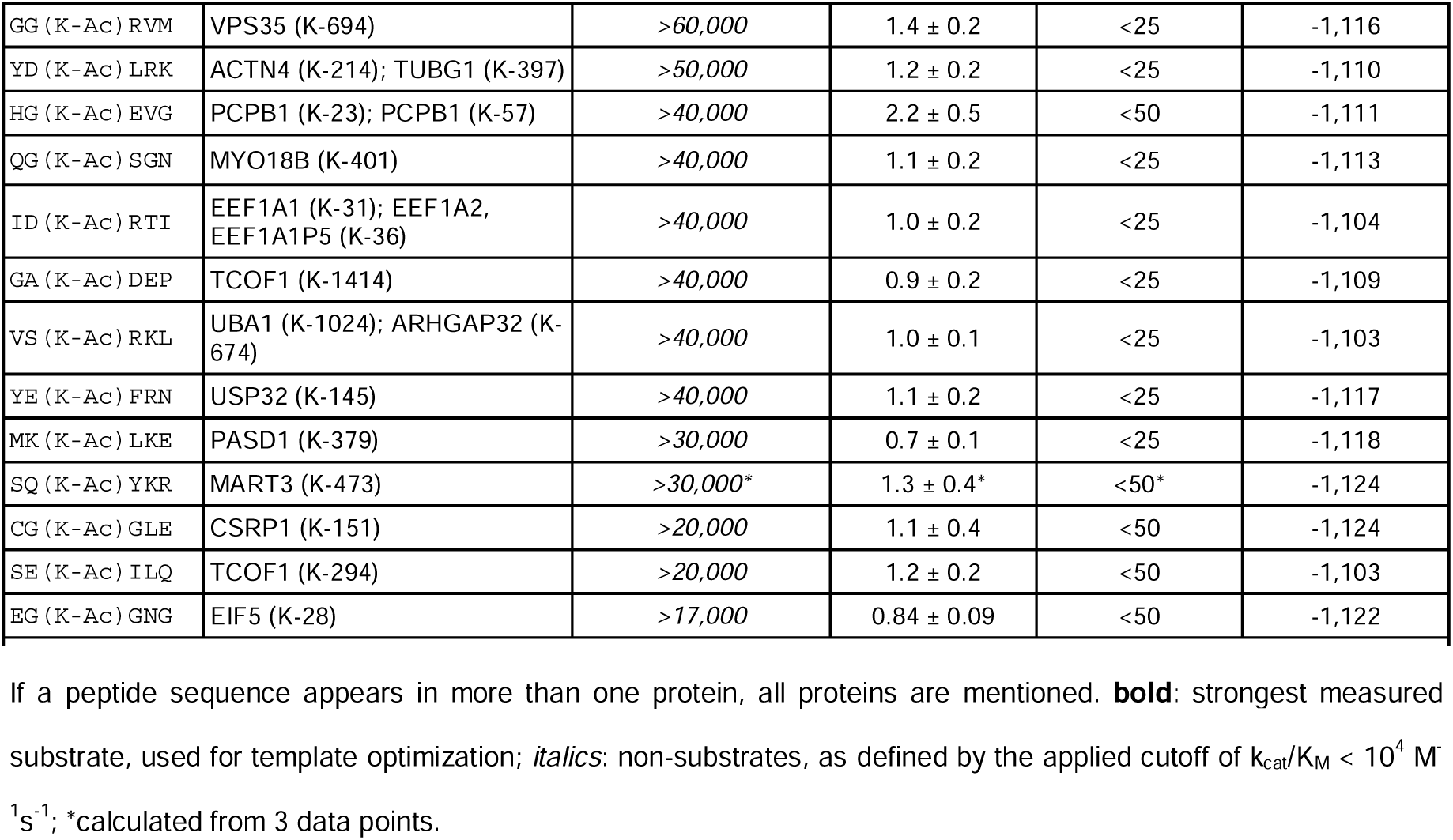
Activities of 42 selected peptides, as measured by the acetate assay. **(A)** Peptide measurements with precise *k*_cat_/*K*_M_ values (D-TRAINING set). **(B)** Peptide substrates with *k*_cat_/*K*_M_ values with lower limits: *K*_M_ values were below the detection limit (D-CAPPED set).

| Peptide | Protein (site of modification) | $k_{\text{cat}}/K_M$<br>($\text{M}^{-1}\text{s}^{-1}$) | $k_{\text{cat}}$<br>($\text{s}^{-1}$) | $K_M$<br>($\mu\text{M}$) | Rosetta<br>reweighted<br>score [REU] |
| --- | --- | --- | --- | --- | --- |
| EG (K-Ac) FVR | LMNA (K-450) | 220,000 ± 40,000 | 4.2 ± 0.7 | 19 ± 4 | -1,123 |
| SD (K-Ac) TIG | TUBA1A, TUBA4A, TUBA3E,<br>TUBA1C, TUBA1B, TUBA4A<br>(K-40)<sup>5,42,43</sup> | 70,000 ± 30,000 | 2.1 ± 0.3 | 29 ± 15 | -1,111 |
| AM (K-Ac) HRS | MYO1G (K-90) | 63,000 ± 9,000 | 1.9 ± 0.2 | 30 ± 7 | -1,114 |
| LA (K-Ac) EMK | CFAP157 (K-35) | 60,000 ± 20,000 | 1.4 ± 0.2 | 20 ± 10 | -1,120 |
| TG (K-Ac) TVA | TCOF1 (K-146) | 50,000 ± 10,000 | 1.37 ± 0.09 | 27 ± 7 | -1,120 |
| RT (K-Ac) SGD | ARHGEF1 (K-234) | 47,000 ± 9,000 | 1.5 ± 0.2 | 32 ± 9 | -1,127 |
| SQ (K-Ac) KTF | GRP94 (K-682) | 44,000 ± 9,000 | 1.6 ± 0.1 | 36 ± 10 | -1,103 |
| AG (K-Ac) RIA | DIP2A (K-962) | 40,000 ± 10,000 | 1.9 ± 0.5 | 50 ± 30 | -1,110 |
| YK (K-Ac) FYE | HSP90A (K-436)<sup>7,8</sup> | 40,000 ± 10,000 | 1.8 ± 0.6 | 21 ± 9 | -1,114 |
| YG (K-Ac) LRK | ACTN1 (K-195) | 31,000 ± 4,000 | 6.0 ± 0.7 | 200 ± 80 | -1,115 |
| EG (K-Ac) TNY | ZNF587 (K-209) | 29,000 ± 7,000 | 2.0 ± 0.4 | 70 ± 30 | -1,110 |
| QK (K-Ac) VKE | ZNF280D (K-32) | 28,000 ± 6,000 | 1.8 ± 0.2 | 60 ± 20 | -1,116 |
| YE (K-Ac) EKE | KIF5A, KIF5C (K-348); KIF5B<br>(K-346) | 27,000 ± 5,000 | 1.8 ± 0.1 | 60 ± 10 | -1,106 |
| EG (K-Ac) TGE | ZKSCAN1 (K-310) | 25,000 ± 4,000 | 1.07 ± 0.08 | 43 ± 9 | -1,112 |
| LS (K-Ac) KSK | PARP1 (K-505) | 24,000 ± 3,000 | 1.7 ± 0.2 | 70 ± 20 | -1,105 |
| PA (K-Ac) ESP | ATBF1 (K-3416); treacle (K-904) | 22,000 ± 4,000 | 1.08 ± 0.05 | 49 ± 9 | -1,112 |
| AM (K-Ac) KIR | CDK1 (K-33) | 16,000 ± 2,000 | 0.98 ± 0.09 | 60 ± 10 | -1,102 |
| QY (K-Ac) KEL | LMNA (K-260) | 13,200 ± 700 | 1.58 ± 0.08 | 120 ± 10 | -1,106 |
| KT (K-Ac) PIW | HSP90A (K-294)<sup>7,8</sup> | 12,000 ± 2,000 | 0.8 ± 0.1 | 60 ± 20 | -1,082 |
| AH (K-Ac) RGS | DPP3 (K-294); CBLL2 (K-107);<br>RFPL4A (K-257) | 9,000 ± 3,000 | >1 | >150 | -1,117 |
| KL (K-Ac) KKE | MYH1 (K-1085) | 5,700 ± 1,100 | 0.50 ± 0.09 | 90 ± 30 | -1,105 |
| EV (K-Ac) KMT | MAP4 (K-847) | 5,600 ± 800 | 0.19 ± 0.02 | 35 ± 7 | -1,105 |
| GY (K-Ac) KTK | RPL4 (K-162) | 5,000 ± 2,000 | 2.3 ± 0.5 | 400 ± 200 | -1,103 |
| AH (K-Ac) KSH | S100A8 (K-84) | 4,700 ± 400 | >0.5 | >100 | -1,099 |
| PL (K-Ac) KDR | RPL3 (K-393) | 2,100 ± 200 | >0.5 | >200 | -1,100 |
| SW (K-Ac) DGL | ACTN2 (K-181) | 1,200 ± 300 | >0.3 | >200 | -1,094 |
| **B** |  |  |  |  |  |
| YD (K-Ac) DEV | SIRT1 (K-430) | >110,000 | 0.28 ± 0.04 | <25 | -1,103 |
| ME (K-Ac) FKI | XPOT (K-627) | >80,000 | 2.0 ± 0.5 | <25 | -1,115 |
| ET (K-Ac) YRW | VDAC1 (K-61) | >70,000 | 3.7 ± 1.2 | <50 | -1,123 |
| GG (K-Ac) RVM | VPS35 (K-694) | >60,000 | 1.4 ± 0.2 | <25 | -1,116 |
| YD (K-Ac) LRK | ACTN4 (K-214); TUBG1 (K-397) | >50,000 | 1.2 ± 0.2 | <25 | -1,110 |
| HG (K-Ac) EVG | PCPB1 (K-23); PCPB1 (K-57) | >40,000 | 2.2 ± 0.5 | <50 | -1,111 |
| QG (K-Ac) SGN | MYO18B (K-401) | >40,000 | 1.1 ± 0.2 | <25 | -1,113 |
| ID (K-Ac) RTI | EEF1A1 (K-31); EEF1A2, EEF1A1P5 (K-36) | >40,000 | 1.0 ± 0.2 | <25 | -1,104 |
| GA (K-Ac) DEP | TCOF1 (K-1414) | >40,000 | 0.9 ± 0.2 | <25 | -1,109 |
| VS (K-Ac) RKL | UBA1 (K-1024); ARHGAP32 (K-674) | >40,000 | 1.0 ± 0.1 | <25 | -1,103 |
| YE (K-Ac) FRN | USP32 (K-145) | >40,000 | 1.1 ± 0.2 | <25 | -1,117 |
| MK (K-Ac) LKE | PASD1 (K-379) | >30,000 | 0.7 ± 0.1 | <25 | -1,118 |
| SQ (K-Ac) YKR | MART3 (K-473) | >30,000* | 1.3 ± 0.4* | <50* | -1,124 |
| CG (K-Ac) GLE | CSRP1 (K-151) | >20,000 | 1.1 ± 0.4 | <50 | -1,124 |
| SE (K-Ac) ILQ | TCOF1 (K-294) | >20,000 | 1.2 ± 0.2 | <50 | -1,103 |
| EG (K-Ac) GNG | EIF5 (K-28) | >17,000 | 0.84 ± 0.09 | <50 | -1,122 |

If a peptide sequence appears in more than one protein, all proteins are mentioned. **bold**: strongest measured substrate, used for template optimization; *italics*: non-substrates, as defined by the applied cutoff of $k_{cat}/K_M < 10^4 \text{ M}^{-1} \text{ s}^{-1}$ ; \*calculated from 3 data points.

**Figure 1.**
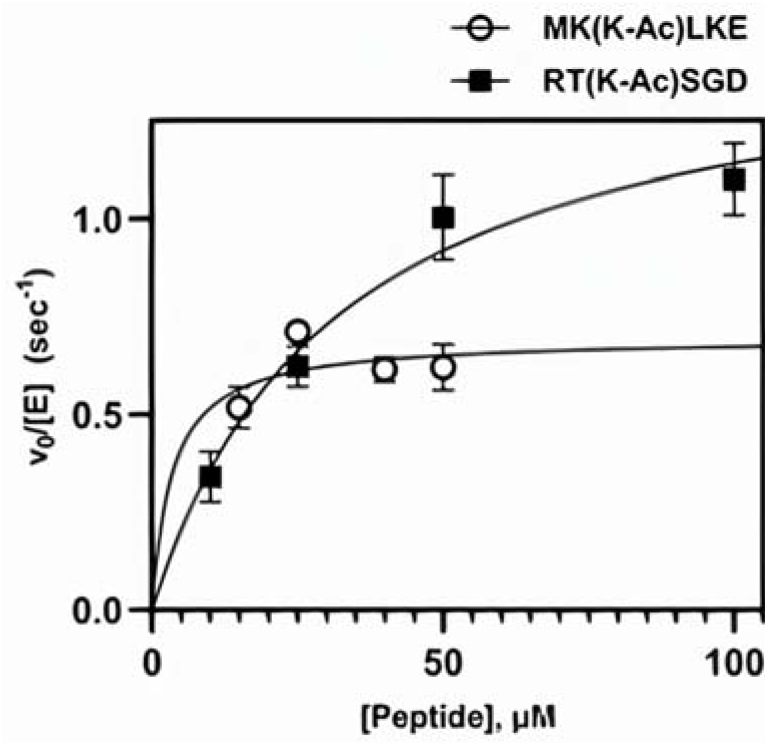
Dependence of deacetylation rate on substrate concentration for two representative peptides catalyzed by HDAC6, measured using the acetate assay. The initial velocity for each substrate concentration was determined from a linear regression of a time course consisting of a minimum of three timepoints, standard error is shown. The kinetic parameters are determined from a nonlinear least square fit of the Michaelis-Menten equation to the data and are listed in **Table 1**. Black squares: example of data which met the criteria to produce accurate *k*_cat_/*K*_M_ values. Open circles: example of overfit data resulting in calculation of a lower limit for *k*_cat_/*K*_M_, due to the *K*_M_ being lower than the detection limit of the assay.

To determine if measured activities on isolated peptides appropriately reflects HDAC6 activity on the natural substrate protein, we measured the activity of HDAC6 towards a full-length, singly-acetylated histone protein and a respective 13-mer peptide analog; catalytic efficiency was similar, albeit higher for the peptide: 14 × 10^4^ M^-1^s^-1^ *vs*. 8 × 10^4^ M^-1^s^-1^ for the full-length substrate, mostly due to an increased value of *k*_cat_. (**Table S2**). Based on these results and the observation that enzymes that are catalytically efficient in the cell have a *k*_cat_/K_M_ value of at least 10^4^ M^-1^s^-1 28^, we defined this value as a cutoff for distinguishing substrate and non-substrate peptides. Additional peptides tested that were not used in further analysis are listed in **Supplementary Table S3** (*i*.*e*., peptides longer than 6 amino acids or peptides that could not be measured reliably).

### Structure-based computational prediction can identify most HDAC6 substrates

To detect new potential substrates for HDAC6, we calibrated a structure-based protocol based on the *k*_cat_/*K*_M_ values for HDAC6-catalyzed deacetylation of hexamer peptides (**Table 1**), using the FlexPepBind framework, previous applied to HDAC8 and FTase enzymes^12,26^. Here we provide a general outline of the calibration process. For details we refer to the Methods section.

First, the HDAC6 substrate with the highest *k*_cat_/*K*_M_ value (EGK_Ac_FVR, derived from prelamin A, see **Table 1**) was docked into the binding pocket of a solved DD2 HDAC6 structure (Protein Data Bank^29^, PDB ID: 6WSJ^30^) using Rosetta FlexPepDock refinement^31^. The top 5 best scoring models were selected as templates. Peptides from the D-TRAINING set were then threaded onto this template set and the peptide-receptor interface was minimized (see Methods). For each peptide, the top-scoring model was used as an estimate for its ability to bind to HDAC6 in a catalysis-competent conformation (reinforced by constraints, see Methods). The performance of the protocol was evaluated based on the calculated binary distinction (area under the curve (AUC) values) and Spearman’s ρ correlation between experimental values and Rosetta scores. The runs were performed with or without receptor backbone minimization, both at the refinement (*ref* vs. *refmin*) and threading steps (*thread* vs. *threadmin*) (*i*.*e*., four different protocols were tested, see summary in **Table 2**). Throughout this study, scoring was performed using the Rosetta reweighted score^32^.

**Table 2.**
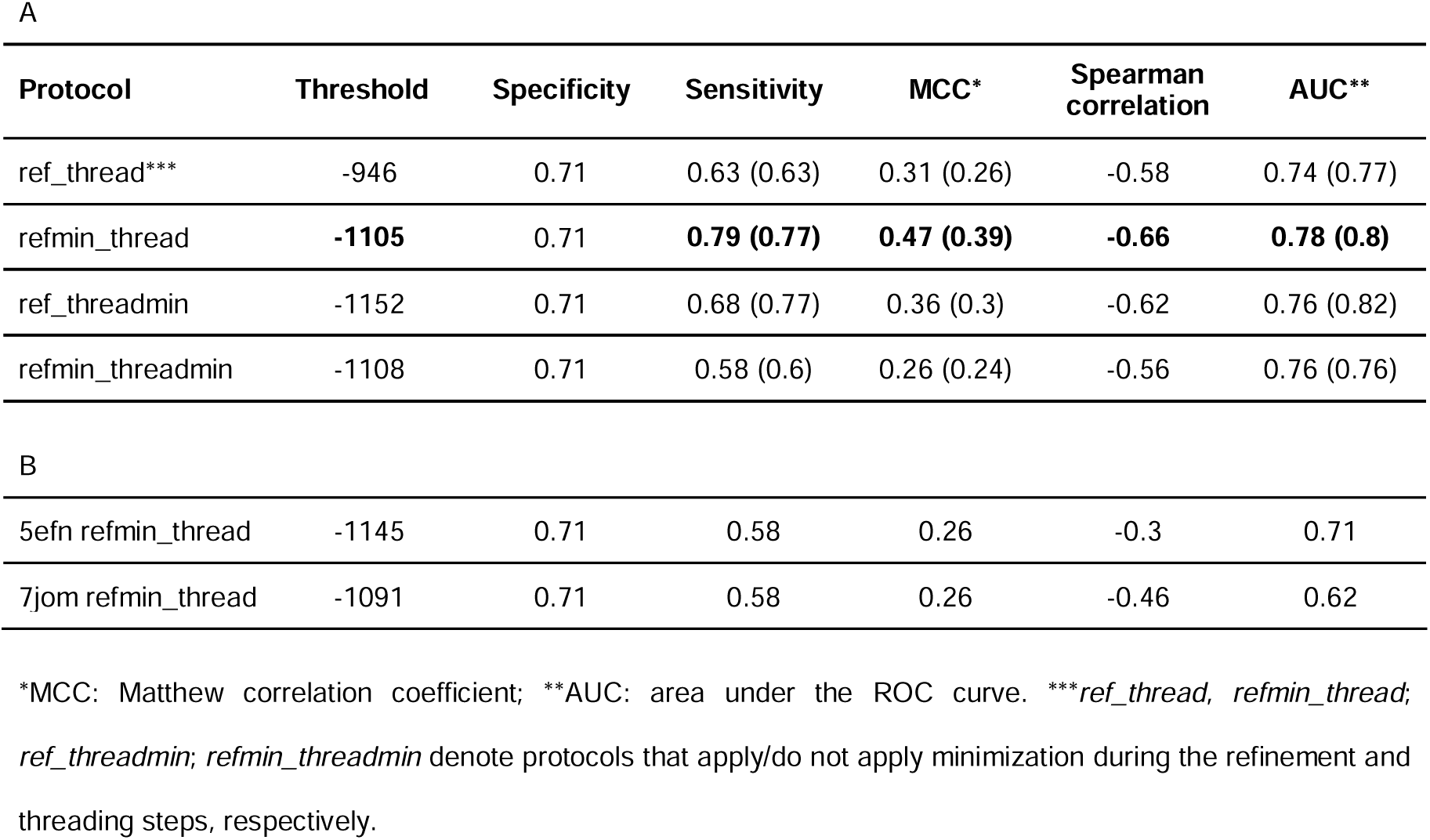
FlexPepBind protocols evaluated in this study. A) Performance metrics of the protocols on the D-TRAINING dataset (n = 26). Values for the dataset including both D-TRAINING and D-CAPPED peptides (n = 26 + 16) are indicated in parentheses. Since the activities of the D-CAPPED dataset are not exact, no correlation was calculated for the merged dataset. Results are shown for the 6WJS template. B) Performance metrics for different crystal structures used as templates, using the *refmin_thread* protocol.

The best performance (AUC = 0.78, ρ = -0.66, **Figure 2**) was achieved when the docking step was performed with backbone minimization, but the subsequent threading step did not include backbone minimization (*i*.*e*., *refmin_thread*). We defined a loose (*reweighted score*: - 1105) and a strict cutoff (*reweighted score*: -1118), to allow for a maximum of 0 and 1 false positives, respectively. Performance was then evaluated on a dataset for which the exact activity values could not be measured but was estimated to consist of substrates only (*i*.*e*., the CAPPED set, see **Table 1**): reassuringly, 12 of the 16 substrates passed the loose threshold (and all 16 passed the strict threshold).

**Figure 2.**
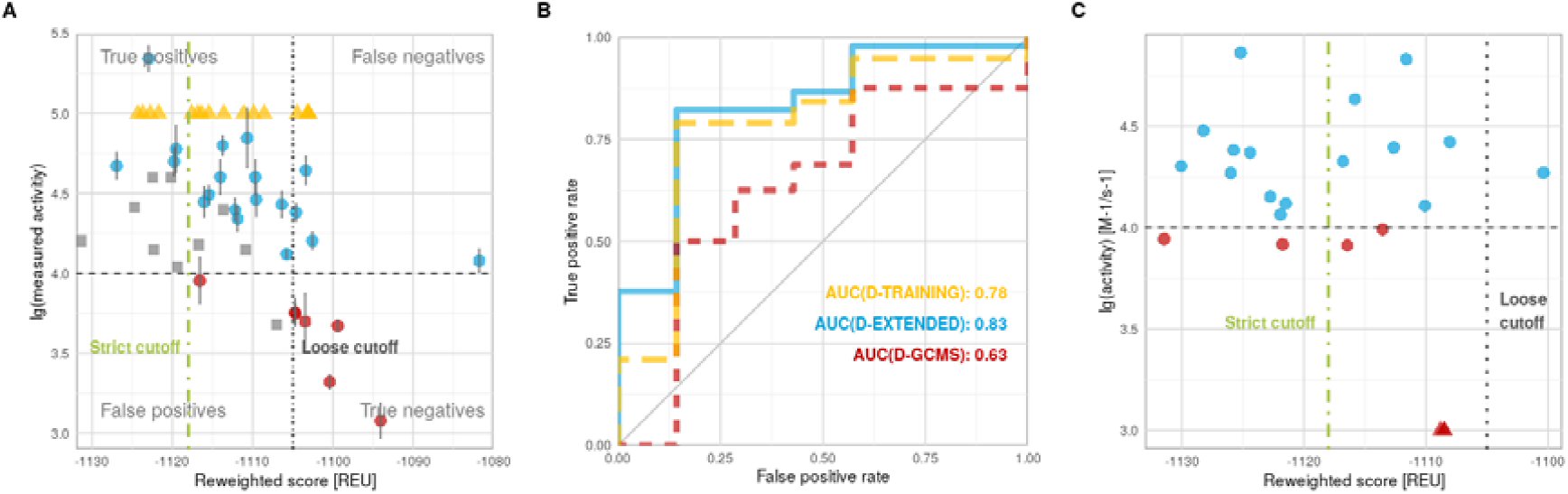
Performance of the calibrated protocol on different datasets (see Supplementary Table S2). **A)** Correlation: Predicted *vs*. measured activities on D-TRAINING (dots, blue: substrates, red: non-substrates); D-CAPPED (yellow triangles); D-TEST (grey squares) datasets. **B)** Binary distinction: ROC curves on D-TRAINING (yellow), D-EXTENDED (blue) and D-GCMS (red). **C)** Performance of protocol on D-GCMS, a dataset measured using GC-MS (from Kutil et al. ^3^). In A and C, dashed horizontal line denotes the cutoff dividing measured substrates and non-substrates (*lg*(*activity*) = 4). Note the different scales in (A) and (C). The cutoffs dividing predicted substrates from not substrates are indicated as dotted vertical line for the loose cutoff (*reweighted scorce* = −1105) and a dotted-dashed vertical line for the strict cutoff (*reweighted scorce*: − 1118).

As independent validation, we applied our protocol to a set of 22 peptides with published measured activities (using GC-MS^3^; the D-GCMS dataset, see **Supplementary Table S1**). Using the same activity cutoff (k_cat_/K_M_ = 10^3^ M^-1^s^-1^), this set is composed of 17 substrates, three non-substrates, and four additional non-substrates with borderline activities (i.e. 10^4^ M^-1^s^-1^ > k_cat_/K_M_ > 8 × 10^3^ M^-1^s^-1^)^3^. Our prediction identified 9 out of 16 substrates using the strict cutoff, and only one was missed by the loose cutoff. The three non-substrates were clearly separated from the rest, lying near the loose cutoff, while the additional four non-substrates with borderline activities showed a range of predicted activities, two passing the stringent threshold and thus predicted to be substrates, i.e. k_cat_/K_M_ = 8.8 × 10^3^ M^-1^s^-1^ and k_cat_/K_M_ = 8.3 × 10^3^ M^-1^s^-1^.

To determine if performance was affected by the specific crystal structure selected we repeated the analysis using different structures. the DD2 domain of HDAC6 was solved several times, bound to different ligands: (1) a cyclic peptide (PDBID: 6WSJ^30^) (2) a tripeptide substrate attached to coumarin (PDBID: 5EFN^4^), and (3) an inhibitor (PDBID: 7JOM^33^). Best performance (shown above) was achieved with the cyclic peptide (6WSJ), while the structure with the inhibitory small molecule performed worst (**Supplementary Figure S1**).

### Predictions on the human acetylome

To detect new potential HDAC6 substrates, we used our calibrated protocol (*refmin_thread* applied to PDB ID 6WSJ) to screen the human acetylome (from PhosphoSitePlus^34^, focusing on peptides annotated from low-throughput experiments). This screen detected 74 peptides that scored better than the peptide with the highest activity in the D-TRAINING set (EGKFVR, *reweighted score*: -1,123, blue line on **Figure 3A**), and 215 and 859 peptides (out of 1,030) were classified as substrates by our strict and loose cutoffs, respectively (belonging to 144 and 297 proteins) (**Figure 3**). In comparison to our previous study on HDAC8 specificity^12^, many more substrates were suggested for HDAC6 (21%) than for HDAC8 (11%) out of the same dataset, which agrees with the reported greater promiscuity observed for HDAC6.

**Figure 3.**
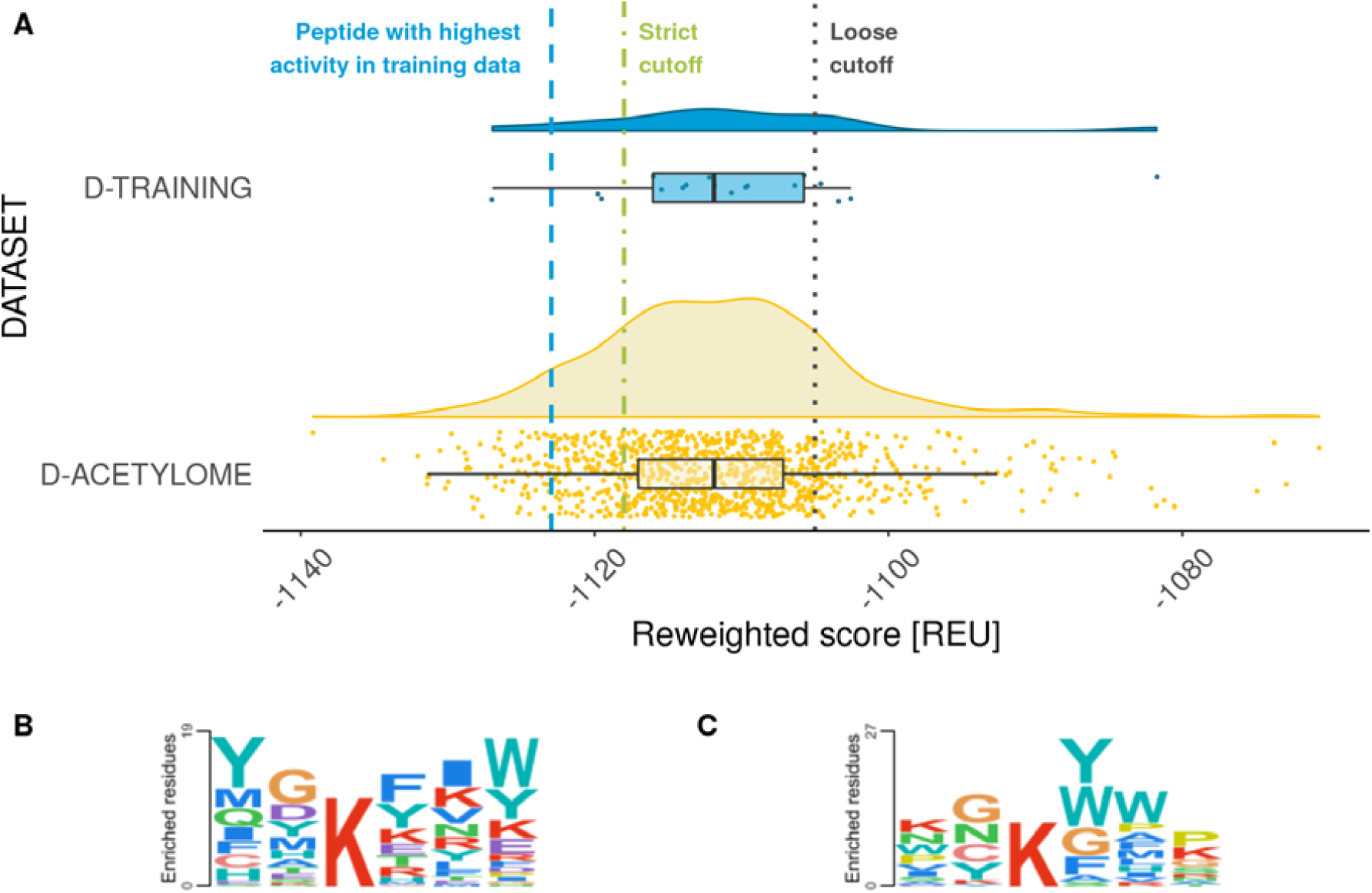
Application of the calibrated protocol to the acetylome to detect novel potential HDAC6 substrates. **(A)** Distribution of scores obtained for acetylated peptides (as annotated in the PhosphoSitePlus database). **(B-C)** Sequence logos of **(B)** substrates from the D-TRAINING set merged with the D-CAPPED dataset, and **(C)** top 100 peptides predicted by our protocol.

Our screen of the known acetylome was able to recapitulate several previously identified substrates, including K49 of β-catenin^35^ (*reweighted score*: -1,129), K274 of the microtubule-associated protein tau 33^36^ (*reweighted score*: -1,120), and K118 of ATP-dependent RNA helicase DDX3X^37^ (*reweighted score*: -1,107). Moreover, we identified 27 proteins that have been previously reported to interact with HDAC6 (based on BioGRID^38^). For these, the interaction could be regulated further by the enzymatic removal of acetylation. In addition to these known substrates and protein interactors, we report several novel hits among the top scored peptides (**Supplementary Data**).

We evaluated the pathway involvement of the proteins associated with the peptide substrates selected for D-TRAINING. Of note, most of the proteins harboring the peptides for which HDAC6 displayed the greatest deacetylase activity (> 4 × 10^4^ M^-1^s^-1^) either play a structural role or are associated with cellular structural elements (LMNA^39^, MYO1G^40^, ACTN1^41^, TUBA1A^5,42,43^), have chaperone functions (GRP94^44^, HSP90^7,8^), or are involved in the Rho signaling pathway (ARHGEF1^45^).

Comparison of the sequence logo created from these peptides to logos created from merged D-TRAINING and D-CAPPED sets (**Figure 3**) reveal that we did not simply recreate the sequence specificity of that dataset. Although there are some similarities, such as the preference for glycine at position P_-1_ and for tyrosine and phenylalanine at P_+1_, the sequence logo is different from the original database and shows more variability. This suggests that this protocol is capable of identifying a broader range of substrates.

### Validating acetylome predictions

From the predicted substrates of the acetylome, we selected 10 peptides for additional validation, using our acetate assay. Out of the 10 peptides, 9 were indeed measured to be substrates (**Table 3**), albeit with poor correlation (ρ = -0.49 for the correctly identified substrates, **Figure 2**). To summarize, our protocol shows robust ability to identify HDAC6 substrates, but due to modest correlation, it’s ability to predict actual substrate strength is limited.

**Table 3.**
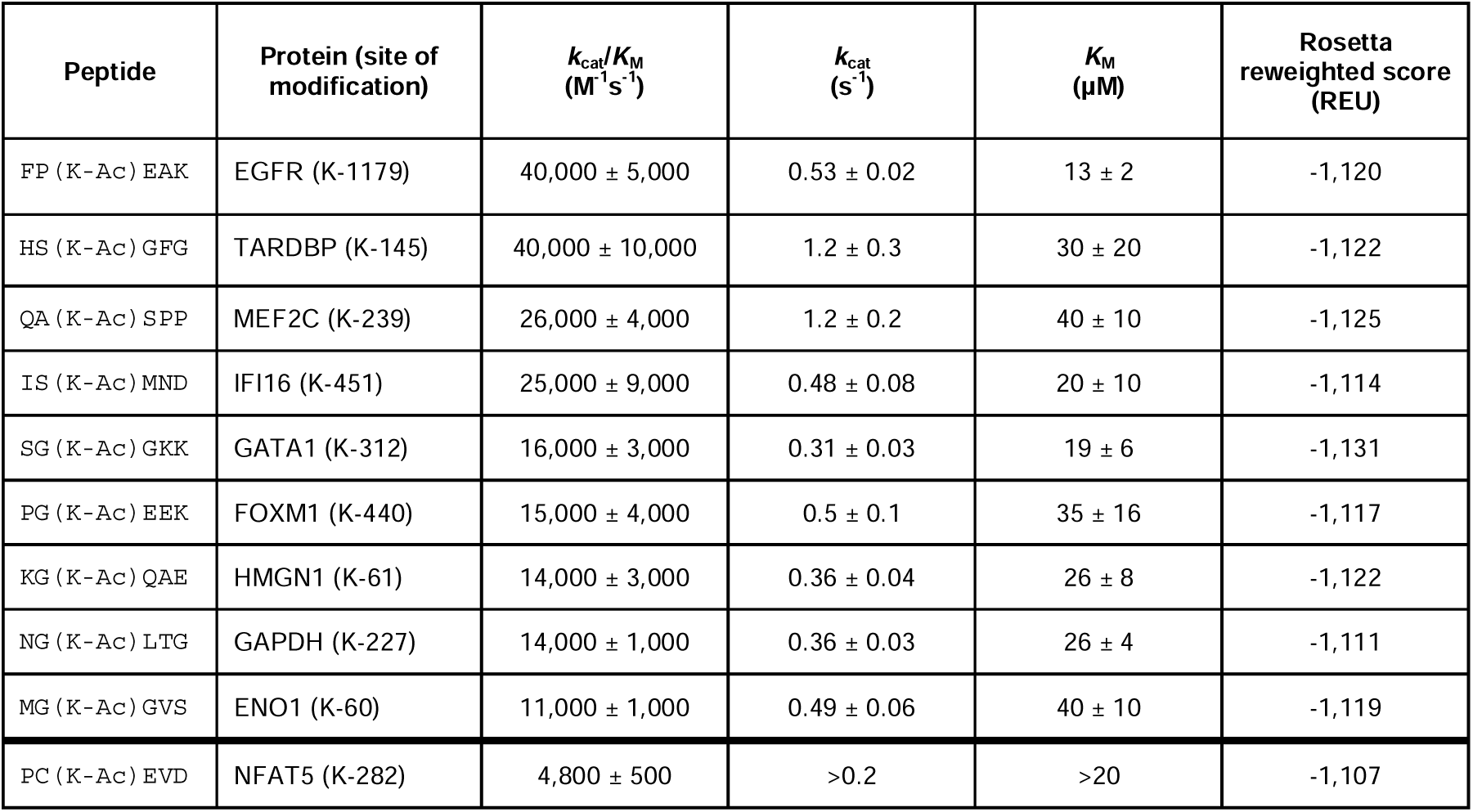
Validation of predicted substrates of the acetylome. The peptides are sorted according to their experimentally measured HDAC6 substrate efficiencies.

### Comparison with previous studies of HDAC6 shows little agreement on substrate selectivity

Several above-mentioned studies have previously probed the substrate landscape to HDAC6. The overlap between substrates at the protein level identified with these different experimental assays was minimal^3^. To further examine these differences, we compared the substrates of these high-throughput methods to the substrates predicted using our protocol. A plot of hexamer peptides shared between D-SILAC and D-13MER, shows poor agreement between the substrate sets of these two studies (**Figure 4**). Consequently, while sequence logos for the respective substrate sets highlight the enrichment of certain residues when compared to a proteome-level (**Figure 4B-D**), or to the non-substrate sets background (**Figure 4E-G)**, they also show significant differences. We used the computed PSSMs (**Supplementary Figure S2**) to cross-score the datasets (using different peptide lengths) and calculated the correlations between the experimental values and PSSM scores (**Supplementary Figure S3**). The strongest, albeit still mediocre correlation (ρ = 0.47) was found between the D-3MER dataset scored with the PSSM derived from itself, and the experimental values of the D-TRAINING dataset scored by the PSSM of the D-13MER or D-SILAC datasets (ρ = 0.46 and 0.44, respectively). This might indicate that different experiments capture different aspects of selectivity and have different biases and limitations. It also means that fitting a computational model considerably better than these experimental agreements would be prone to overfitting.

**Figure 4.**
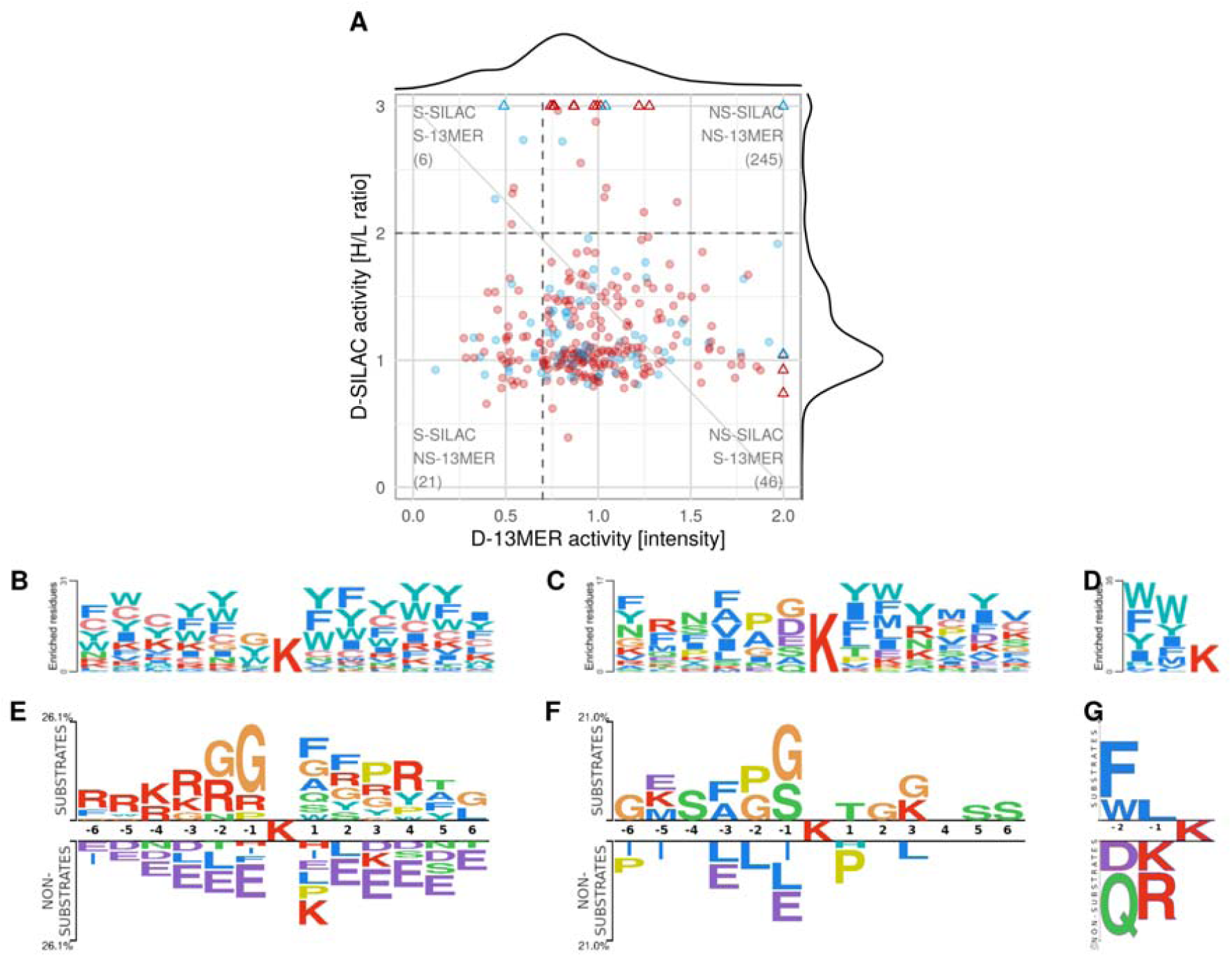
Different datasets of experimentally determined HDAC6 substrates show minor overlap and agreement. Comparison of substrate specificities of different experimental datasets, based on their sequence logos and correlation of substrate activities. **(A)** Plot of substrate activities for peptides shared between D-13MER ^3^ and D-SILAC ^14^ datasets. **(B-F)** Sequence logos made with PSSMSearch using its default background containing human sequences (B-D), and with Two Sample Logo using the non-substrate peptides as background for each dataset (E-G). **(B**,**E)** D-13MER, **(C**,**F)** D-SILAC, and **(D**,**G)** D-3MER datasets. S, NS: substrates and non-substrates, respectively.

## Discussion

In this study we have calibrated a structure-based protocol to characterize HDAC6 substrates, and applied this protocol to identify new potential deacetylation substrates. We trained the method on a set of selected peptides for which we measured catalytic activity, validated the model on a set of independently measured peptides and applied it to the human acetylome. In the following discussion we compare the selectivity determinants of HDAC6 and HDAC8, summarize potential roles for the newly predicted substrates, and point to challenges in the accurate and comprehensive characterization of a promiscuous enzyme such as HDAC6.

### Differences between HDAC6 and HDAC8

HDAC6 has a much higher catalytic efficiency and increased promiscuity towards peptide substrates *in vitro* compared to that of HDAC8 reported previously. For example, the fastest measured peptide substrate of HDAC8, ZNF318 at K1275, has a *k*_cat_/*K*_M_ value of 4.8 × 10^3^ M^-1^s^-1 12^ (over 40-fold lower than the fastest HDAC6 substrate; it would not be considered a substrate for HDAC6). It is known that the activity of HDAC8 towards peptides is much lower than its activity towards full-length protein substrates^46^. However, this is not the case for HDAC6, at least DD2, as activity of peptides and full-length protein substrates are similar (**Supplementary Table S2**) where HDAC8 sees an increase in catalytic efficiency of about 100-fold, HDAC6 displays catalytic efficiencies within a 2-fold range. Similarity between activity towards short peptides and full-length protein substrates indicates that HDAC6 substrate preference is determined through short range interactions between the active site and the substrate, and peptides should be an appropriate analog for full-length substrate activity. We further found that the increased catalytic efficiency was linked to the low *K*_M_ values of most of the peptides. To develop our protocol, we determined it was necessary to obtain accurate measurements of catalytic efficiency, as the low *K*_M_ values easily allow the *k*_cat_/*K*_M_ value to be underestimated using measurements with a single substrate concentration.

Even with the assurance of accurate activity measurements, predicting the selectivity of HDAC6 proved to be much more challenging than its paralog HDAC8 for which we were able to obtain good predictions without introducing any backbone receptor flexibility^12^. We explored structural differences between these proteins that could explain this finding. Comparison of the HDAC6 (PDB ID: 6WSJ^30^) and HDAC8 (PDB ID: 2V5W^47^) structures (**Figure 5**) highlight the main difference is in a loop involved in forming the binding pocket that could lead to differences in binding selectivity. The residue that forms a hydrogen bond with the acetylated lysine to position the substrate, D101 in HDAC8 and S531 in HDAC6, contacts the backbone at a similar position, but stems from a very different loop backbone. This allows for the downstream part of the peptide to make larger movements, accommodating more substrates. The loop also participates in the formation of the pocket accommodating the residue preceding the acetylated lysine (P_-1_). Indeed, in HDAC6 this pocket is considerably smaller, explaining the significant enrichment for glycine in the peptide libraries. This loop in HDAC8 also harbors Y100, a residue whose hydroxyl group forms a hydrogen bond with the peptide backbone in HDAC8, providing an additional recognition feature. No residue corresponding to Y100 is found in HDAC6.

**Figure 5.**
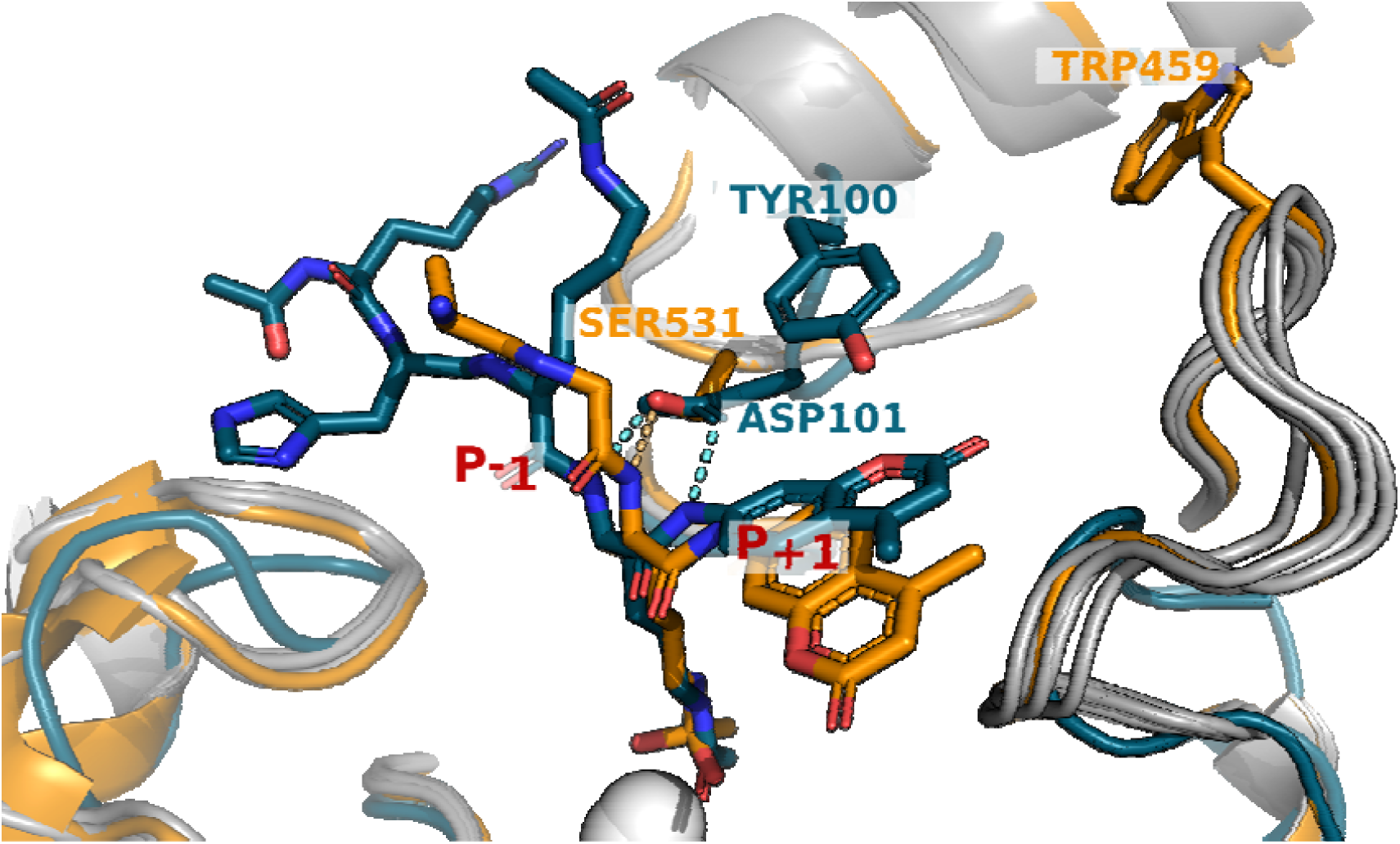
Key differences between HDAC6 and HDAC8 in the coordination of the substrate. The active sites of HDAC6 (receptor from PDB ID 6WSJ, substrate from 5EFN for visualization purposes, orange) and HDAC8 (PDB ID 2V5W, teal) are overlaid. The loops that distinguish the two HDACs are highlighted (orange and teal), together with conformations generated for modeling substrate activity (grey). Residues S531 (HDAC6) and D101 (HDAC8) that coordinate hydrogen bonding (light blue: HDAC6, yellow: HDAC8) to the backbone of the acetylated substrate lysine residue, are also highlighted, as well as positions W459 (HDAC6) and Y100 (HDAC8). The hydroxamic acid inhibitors are shown in sticks, and the catalytic Zn^2+^ ions are shown as a white sphere. See Text for more details.

The loop located near the pocket that accommodates the residues trailing the acetylated lysine (P_+1-3_) is significantly longer (12 residues in HDAC6 compared to 7 in HDAC8) and more hydrophobic in HDAC6. The larger size of the loop, together with its more non-polar character, suggest that the HDAC6 loop may be more flexible, allowing for adaptations of the binding groove, resulting in a more promiscuous binding pattern. Indeed, we showed here that receptor backbone minimization, that moves this loop, is needed for our protocol to succeed, suggesting that this pocket is restructured for binding.

Previous attempts for modeling the interactions between the measured peptides only gave mediocre results, although multiple starting structures from PDB were evaluated (e.g. 5EFN, 5EFK, 5EDU, etc, data not shown). These structures were either bound to small molecule inhibitors or tripeptide substrates covalently linked to 7-amino-4-methylcoumarin. However, with the release of 6WSJ, HDAC6 DD2 bound to a cyclic peptide, the predictions improved. Evaluation of 3 starting structures with different entities in the binding pocket (5EFN - tripeptide substrate linked to coumarine^4^, 6WSJ and 7JOM - small molecule inhibitor^33^) reveal the best predictive power using DD2 bound to a cyclic peptide (6WSJ) and the worst belonging to DD2 in complex with a small molecule (See **Supplementary Figure S1** and **Table 2B**). This highlights the importance of choosing the right starting structure.

### Impact of predicted substrate deacetylation

Our substrate prediction model suggests novel aspects of HDAC6 deacetylase function. Acetylation and ubiquitination reportedly compete for lysine residues; acetylation can prevent lysine ubiquitination^48,49^ or ubiquitin chain elongation^50^ and consequently protect against proteasomal degradation. We inspected the acetylated lysines in our predicted substrates, as well as their flanking regions, for additional reported post-translational modifications. Among the 215 candidates that passed the strict score cutoff of -1118, 93 (43%) underwent ubiquitination (data from PhosphoSitePlus^51^, see **Supplementary Data**). Furthermore, deacetylation of some of these sites has already been linked to promoting protein ubiquitination and degradation: K569 and K259 of Forkhead box protein O3 (FOXO3^52^), K709 of Hypoxia inducible factor 1 alpha (HIF1A^53^), K887 and K1413 of Werner syndrome ATP-dependent helicase (WRN^54^), K406 of Chorion-specific transcription factor GCMa (GCM1^55^), and K540 of ATP-citrate synthase (ACLY^56^). Additional exploration of the role of HDAC6, deacetylation, and ubiquitination could illuminate important regulation of the involved biological pathways.

### Novel potential regulatory functions of HDAC6

The success of the substrate prediction model is shown in its ability to reinforce previously identified HDAC6 substrates as well as allow novel HDAC6 functions to be explored. The two measured acetylome peptides with the best HDAC6-catalyzed deacetylation values (4 × 10^4^ M^-1^s^-1^), are derived from the proteins of EGFR and TARDBP. TARDBP has been previously explored as a direct substrate of HDAC6: acetylated TARDBP aggregates are found in patients with amyotrophic lateral sclerosis (ALS)^57^ and deacetylation of TARDBP by HDAC6 prevents TARDBP aggregation^57^.

HDAC6 is known to regulate EGFR endocytic trafficking between the apical and basolateral membranes by regulating deacetylation of α-tubulin and affecting its turnover rate^58,59^. Acetylation of EGFR has been shown to enhance its activity, an effect observed through treatment with HDAC inhibitors^60^. Two high-scoring peptides harboring residue K1179 and K1188 (*reweighted scores*: -1120 and -1122, respectively) are derived from EGFR. We hypothesize that HDAC6 could play a role in the regulation of the protein level and activity not just by catalyzing deacetylation of EGFR interactors (i.e., α-tubulin), but also by deacetylating EGFR itself.

Reactome Pathway analysis^61^ of the proteins belonging to the top 100 best scoring peptides identified pathways of *Estrogen-dependent gene expression* (R-HSA-9018519, FDR=0.01) and *Transcriptional regulation of granulopoiesis* (R-HSA-9616222, FDR=0.03) as significantly overrepresented compared to the background of the human proteome (FDR<0.05). However, the latter, although significant, was also enriched when the whole acetylome was compared to the background. Several studies have shown a link between HDAC6 and estrogen-signaling^62,63^, with estrogen upregulating the level of HDAC6^62,63^, and we also identified the estrogen receptor as a possible substrate (K171, *reweighted score: - 1119*).

One of our strongest predicted substrate peptides harbors K87 on nucleolin (NCL, *reweighted score:* -1130) which is predicted^16^ to be part of a cyclin-dependent kinase (CDK) phosphorylation motif (MOD_CDK_SPxK_1 for K87 or MOD_CDK_SPxxK_3 for K88), where the charge of the lysine is crucial for recognition^64^. Therefore phosphorylation would probably be hindered by acetylation of the lysine residue. Phosphorylation of T84 in NCL was shown to be crucial for its interaction with DEAD box polypeptide 31 (DDX31)^65^. This complex is important for activating EGFR/Akt signaling, which induces cell survival and proliferation^66^ and Akt was shown to be hypo-phosphorylated in HDAC6 knockout mice^67^. This could provide further explanation for the role of HDAC6 in tumorigenesis and tumor survival^59,67^, pointing to a multi-level regulation of these processes.

Some of the acetylated sites (K1699, K1550, K1769, K1546) in histone acetyltransferase p300 (EP300) are predicted to be putative substrates. Although the deacetylation of EP300 was previously linked to several other HDACs^48,49^, these data suggest that HDAC6 could also be a potential regulator of this protein. This reveals an intriguing possibility of mutual crosstalk via PTMs, since EP300 has been shown to acetylate HDAC6 and thereby modify the ability of HDAC6 to deacetylate tubulin^50^.

Several proteins identified in the acetylome screen are involved in the cellular response to oxidative stress. These proteins are transcription factors which function through intracellular localization changes in response to cellular signaling, particularly nuclear factor erythroid 2-related factor 2 (NRF2), HIF1A, FOXO3, and Forkhead box protein O1 (FOXO1). The acetylation of transcription factor FOXO1 has been connected to the protein residing in the cytoplasm, and deacetylation results in nuclear importation/retention to allow for DNA binding and transcriptional activation^68^. The acetylome screen identified several of FOXO1 acetylation sites as potential HDAC6 substrates.

The acetylome screen also revealed numerous predicted substrates involved in metabolism with important links to cancer development. Our second best scoring substrate, K311 of Glutaminase (*reweighted score*: -1134, GLS), plays an important part in regulating GLS oligomerization which is crucial for the activation of the enzyme^69^. Removal of the acetyl group induces oligomerization and activation of GLS which reduces oxidative stress in cancer cells therefore promoting their survival^70^. Glycolytic enzymes have been proposed to be substrates of HDAC6^71^, and our acetylome screen identified two glycolytic enzymes shown to interact with HDAC6, phosphoglycerate kinase 1 (PGK1)^72^ and pyruvate kinase muscle (PKM)^73^. The acetylome screen identified additional metabolic enzymes, including heme oxygenase 1 (HMOX1)^74^, ATP-citrate lyase (ACLY)^56^, and malic enzyme 1 (ME1)^75^. The acetylation of such enzymes has been connected to regulation of enzymatic activity leading to carcinogenesis. Our results indicate the involvement of HDAC6 in cancer may be more related to metabolic pathways than previously proposed.

### Comparison to previous HDAC6 substrate studies

In this study we develop a structure-based model of HDAC6 substrate selectivity, and successfully validate it experimentally on a number of substrates. Nevertheless, comparison to other published studies shows limited agreement of detected HDAC6 substrates among any of these studies (**Figure 4**). This disagreement could be due to different experimental setups. One potential difference is using only the DD2 domain of HDAC6 instead of the full-length protein. However, the studies discussed here (D-3MER^13^, D-SILAC^3^ and D-13MER^14^ datasets) all used the full-length enzyme and still report little overlap (e.g., only 5 common substrates between D-13MER and D-SILAC). Another source of variation could stem from the varied size of the peptides and their flanking regions in the different experiments. Among the peptides measured both in the D-13MER and D-SILAC datasets, there were 171 and 89, respectively, whose core hexamers were the same, but their flanking regions differed. Nevertheless, most were either substrates or non-substrates irrespective of the different flanking regions (**Supplementary Figure S4**). Closer inspection suggests that most of these variations could be explained by experimental measurement variations (*i*.*e*., outliers among the repeated experiments in case of D-13MER and protein inference differences for the D-SILAC dataset. As for the latter, assigning peptides with overlapping sequences to different proteins affects their quantification). It therefore does not come as a surprise that PSSMs generated based on the different datasets show poor resemblance (**Figure 4** and **Supplementary Figure S2**).

None of these PSSM matrices could be used to generate predictions that correlate strongly with experimentally measured values (best correlation R=0.47, see **Supplementary Figure S3**). Of note, PSSMs treat peptide positions independently and do not incorporate additional information, e.g. secondary structure and disorder, thus may not be able to fully encompass substrate selectivity determinants. Overall this suggests that due to possible experimental biases, the different experiments only capture part of the specificity, and consequently would also explain our suboptimal performance on these datasets.

*In vivo* experiments evaluate HDAC6 substrate selectivity in its physiological context, including potential cofactors, binding proteins and post-translational modifications of both the substrates and HDAC6. For example, HDAC6 displays different affinities toward tubulin dimers compared to assembled microtubules^76^. In *in vitro* experiments, peptides outside of their native environment might not fold into their native secondary structure that may be important for recognition^77^. Moreover, *in vivo* experiments can also detect downstream effects of treatments, therefore introducing bias into the results. Additionally, as with all high throughput experiments, they have a higher chance of amplifying noise. Furthermore, antibodies widely used to detect or enrich for acetylation might not be sensitive or specific enough or have a bias for certain amino acids around the modification site^78^, and mass spectrometry is biased to capture peptides where the positive charge is not removed by post-translational modifications such as acetylation^79^.

## Conclusion

The present study provides for the first time accurately measured HDAC6 enzymatic activities on a large set of unlabeled peptides. The prediction model developed here allowed for the discovery of novel substrates and novel avenues of substrate exploration for HDAC6. The construction of a structure-based model for a second HDAC isozyme highlights the utility and limitations of this approach for predicting novel substrates for isozymes with very different substrate ranges. The promiscuity of HDAC6 is particularly apparent in the limited agreement amongst the different large-scale substrate selectivity datasets. Our prediction model illuminated possible structural determinants of the broad selectivity of HDAC6 in comparison to HDAC8 and suggested novel regulatory roles for this enzyme.

## Materials and Methods

### Measuring enzyme activity

#### Reagents

High flow amylose resin was purchased from New England Biolabs and Ni-NTA agarose was purchased from Qiagen. Adenosine triphosphate (ATP), coenzyme A (CoA), NAD^+^, NADH, L-malic acid, malate dehydrogenase (MDH), citrate synthase (CS), and mouse monoclonal anti-polyhistidine-alkaline phosphatase antibody were purchased from Sigma. Monoacetylated peptides, with N-terminal acetylation and C-terminal amidation, were purchased from Peptide 2.0 or Synthetic Biomolecules. 3% (v/v) acetic acid standard was purchased from RICCA Chemical. All other materials were purchased from Fisher at >95% purity unless noted otherwise.

### HDAC6 Expression and Purification

The plasmid and protocol for the expression and purification of *Danio rerio* HDAC6 catalytic domain 2 (DD2) was generously provided by David Christianson (University of Pennsylvania). The expression construct was prepared previously by the Christianson lab by cloning the residues 440-798 of the *Danio rerio* HDAC6 gene into a modified pET28a(+) vector in frame with a TEV-protease cleavable N-terminal 6xHis-maltose binding protein (MBP) tag^4^. HDAC6 was expressed and purified as described with several alterations for expression optimization ^4^. BL21(DE3) *E. coli* cells (Novagen 69450-3) were transformed with plasmid according to the protocol and plated on LB media-agar supplemented with 50 μg/mL kanamycin. Plates were incubated overnight at 37°C (16-18 hours), and a single colony was added to a LB media starter culture supplemented with 50 μg/mL kanamycin and incubated with shaking at 37°C for 16-18 hours. This overnight starter culture was diluted (1:200) into 2x-YT media supplemented with 50 μg/mL kanamycin and incubated at 37°C with shaking until the cell density reached an OD600=1. The cultures were then cooled to 18°C for one hour and supplemented with 100 μM ZnSO4 and 500 μM isopropyl β-D-1-thiogalactopyranoside (IPTG) to induce expression. The cultures were grown for an additional 16-18 hours with shaking at 18°C and harvested by centrifugation at 6,000 x g for 15 min at 4°C. Cell pellets were stored at -80°C. 1-mL pre- and post-induction samples were taken and tested for HDAC6 expression by polyhistidine western blot and activity using the commercial Fluor de Lys assay (Enzo Life Sciences).

Cell pellets were resuspended in running buffer (50 mM HEPES, pH 7.5, 300 mM KCl, 10% (v/v) glycerol and 1 mM TCEP) supplemented with protease inhibitor tablets (Pierce) at 2 mL/g cell pellet. The cells were lysed by three passages through a chilled microfluidizer (Microfluidics) and centrifuged for 1 h at 26,000 x g at 4°C. Using an AKTA Pure FPLC (GE) running at 2 mL/min, the cleared lysate was loaded onto a 10-mL packed Ni-NTA column equilibrated with running buffer. The column was washed with 10 column volumes (CVs) of running buffer and 10 CVs of running buffer containing 30 mM imidazole, and the protein was eluted with 5 CVs elution buffer containing 500 mM imidazole. 8 mL fractions were collected and analyzed by SDS-PAGE and western blot, and fractions containing His-tagged HDAC6 were combined and loaded onto a 30-mL amylose column equilibrated with running buffer at 1 mL/min. The column was washed with 2 CVs running buffer and the protein was eluted with 5 CVs of running buffer supplemented with 20 mM maltose. Fractions containing HDAC6 were combined with His6x-TEV S219V protease (0.5 mg TEV protease/L culture), previously purified in-house ^80^ using a commercially purchased plasmid (Addgene plasmid pRK739), and dialyzed in 20K molecular weight cut-off (MWCO) dialysis cassettes against 200-fold running buffer containing 20 mM imidazole overnight at 4°C. After dialysis, the sample was loaded onto a 10-mL Ni-NTA column pre-equilibrated with running buffer containing 50 mM imidazole at 2 mL/min. The column was washed with 5 CVs of 50 mM imidazole running buffer to elute cleaved HDAC. Non-cleaved HDAC6 and His-tagged TEV-protease were eluted with 20 CVs of a 50-500 mM linear imidazole gradient. Fractions containing cleaved HDAC6 were combined, concentrated to <2 mL, and loaded onto a 26/60 Sephacryl S200 size exclusion chromatography (SEC) column (GE) equilibrated with SEC/storage buffer (50 mM HEPES, pH 7.5, 100 mM KCl, 5% glycerol, and 1 mM TCEP) at 0.5 mL/min. Eluted peaks were tested for deacetylase activity, and active fractions were concentrated, aliquoted, flash frozen with liquid nitrogen, and stored at -80°C.

### Acetyl-CoA Synthetase (ACS) Expression and Purification

The pHD4-ACS-TEV-His6x expression vector was prepared previously by inserting the ACS gene from a chitin-tagged acetyl-CoA synthetase plasmid Acs/pTYB1, a generous gift from Andrew Gulick (Hauptman-Woodward Institute), into a pET vector containing a His6x tag to increase expression^12,27^. The pHD4-ACS-TEV-His6x construct was expressed and purified as previously described^27^.

### Coupled Acetate Detection Assay

The coupled acetate detection assay or simply the ‘acetate assay’ was performed as previously described with a few modifications^27^. Briefly, lyophilized peptides were re-suspended in water when possible or with minimal quantities of acid, base, or organic solvent to improve solubility. Peptide concentration was determined by one or more of the following methods: 1) measuring A_280_ using the extinction coefficients if the peptide contained a tryptophan or tyrosine, using the fluorescamine assay if the peptide contained a free lysine^81^, 2) performing the bicinchoninic (BCA) assay using bovine serum albumin (BSA) as a standard, and 3) determining the concentration of acetate produced by complete deacetylation of the peptide by HDAC6.

Reactions containing 10-2000 μM monoacetylated peptides in 1X HDAC6 assay buffer (50 mM HEPES, pH 8.0, 137 mM NaCl, 2.7 mM KCl, 1 mM MgCl_2_) were initiated with 0.1-1 μM HDAC6 at 30°C. Timepoints, 60 μL, were quenched with 5 μL of 10% hydrochloric acid and kept on ice until assay completion (no more than 90 minutes). Timepoints were flash frozen with liquid nitrogen and stored at -80°C until work-up.

Coupled solution (50 mM HEPES, pH 8, 400 μM ATP, 10 μM NAD^+^, 30 μM CoA, 0.07 U/μL CS, 0.04 U/μL MDH, 50 μM ACS, 100 mM NaCl, 3 mM KCl, 50 mM MgCl_2_, and 2.5 mM L-malic acid) was prepared the day of the work-up and incubated at room temperature away from light for at least 25 minutes. Timepoints were quickly thawed and neutralized with 15 μL of freshly prepared and filtered 6% sodium bicarbonate. Neutralized timepoints or acetate or NADH standards, 60 μL, were added to 10 μL coupled solution (or 1X assay buffer for NADH standards) in a black, flat-bottomed, half-area, non-binding, 96-well plate (Corning No. 3686). The resulting NADH fluorescence (E_x_=340 nm, E_m_=460 nm) of standards and timepoints was read on a PolarStar fluorescence plate reader at 1-3-minute increments until the signal reached equilibrium. The acetate and NADH standard curves were compared to verify the coupled solution’s activity. When possible, a positive control reaction for enzyme activity was included. Using the acetate standard curve, the fluorescence of each timepoint was converted to μM product, and the slopes of the linear portion of the reaction (<10%) were plotted against substrate concentration. Using GraphPad Prism, the Michaelis-Menten equation (**Equation 1**) was fit to the resulting dependence of the initial velocity on substrate concentration to determine the kinetic parameters *k*_cat_/*K*_M_, *k*_cat_, and *K*_M_. Standard error was calculated using GraphPad Prism analysis. A total of 50 peptides were tested. Of those, 26 were used in the training set and another 16 were used for validation of the protocol (**Table 1, Supplementary Table S1**). 8 of the peptides were longer than 6 residues or their activity could not be measured precisely, therefore we did not use them for further analysis .

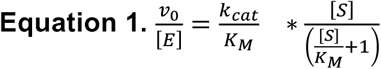

### Calibration of FlexPepBind

The protocol implemented in this study is similar to the one used in our previous study on HDAC8 specificity^12^, and in the following we mainly highlight the differences. In every Rosetta protocol described, we used Rosetta v2020.28. See **Supplementary Table S6** and **Supplementary Table S7** for command-line files and arguments for running FlexPepDock.

Running FlexPepBind requires the creation of a starting structure to generate a template (or a set of templates, as in the present study) for threading peptides. We used the structure of the *Danio rerio* HDAC6 catalytic domain 2 (DD2) which was crystallized in a complex with a cyclic peptide substrate (PDB ID: 6WSJ^30^). To enforce a catalysis-competent binding conformation, we defined constraints that characterize substrate binding as revealed by the solved structures of HDAC6 bound to ligands. Constraints were defined for Rosetta runs as with HDAC8^12^. These include: (1) interactions coordinating the proper binding of the Zn^2+^ ion required for enzymatic activity, (2) interactions between the acetylated lysine side chain and the binding pocket, (3) a dihedral angle constraint in the peptide between residues 3 and 4 (*i*.*e*., adjacent to the acetylated lysine in the modeled hexamers) to enforce a cis-peptide bond (see **Supplementary Table S6** for further details). All of the distances between interacting residues were measured on structure PDB ID 6WSJ. For comparing the final protocol on structures 5EFN, 6WSJ and 7JOM, the constraints were measured on the respective structures.

The best substrate peptide (sequence: *EGK*_*Ac*_*FVR*) was built into the binding pocket of 5EFN, using the trimer substrates and corresponding atoms of coumarin, with the trailing 3 residues of the peptide being added in extended conformation. Then, the resulting complex was superimposed onto 6WSJ and the peptide conformation copied into its binding pocket. FlexPepDock was run on this structure with the constraints added, generating *nstruct*=250 decoys with different setups (**Table 2**), both with and without receptor backbone minimization (*refmin* and *ref* protocols, respectively). The scoring function of ref2015^82^ was used and Reweighted score (reweighted_sc) was found to best discriminate substrates (compared to total score and interface score). Reweighted score is calculated by summing Rosetta’s total score, interface score and peptide score, giving double weight to interface residues and triple weight to the residues of the peptide. In contrast to the HDAC8 study, we selected not only the top-scoring structure, but rather the top 5 structures, according to *reweighted_sc*. Every peptide of the training dataset was threaded onto these starting structures using the Rosetta *fixbb* protocol and running FlexPepDock with minimization only (see **Supplementary Table S6** and **Supplementary Table S7** for runline commands and parameters). Again, these simulations were run with or without receptor backbone minimization (*threadmin* and *thread*, respectively), resulting in four different potential protocols: *refmin_threadmin, refmin_thread, ref_threadmin*, and *ref_thread*. For each peptide sequence, the best score (*reweighted_sc)* among the 5 templates was used to reflect its substrate strength.

### Comparison of datasets

The PSSMs for the comparison of different datasets were generated with PSSMSearch^83^, using PSI-BLAST as the scoring method. Substrates were defined according to the thresholds applied in the reported studies, except for the D-3MER dataset, where no such threshold was provided. The substrates for this dataset were taken from the first five out of a total of 15 bins. The differential sequence logos were generated with the TwoSampleLogo standalone program^84^, using default values. Correlation and AUC calculations and visualizations were created in R (v3.5.1), using *corrplot* (version 0.84) and *pROC* (v1.16.2^85^) packages, respectively.

For the calculation of the scores using the PSSM of the D-3MER dataset, we used only the leading two amino acids before the acetylated lysine in the substrate. We also scored the peptides by only using values from D-SILAC and D-13MER PSSMs in P_-1_ and P_-2_ positions.

In summary, every 3-mer peptide has three different scores and every 13-mer peptide has 7 different scores (4 for the full-length peptide based on D-SILAC, and D-13MER with 13 and 6 scored amino acids; and 3 for assessing the core 3-mers).

Common peptides from the D-13MER and D-SILAC dataset were selected for comparison of the two approaches. For defining substrates and non-substrates, we used the thresholds of 0.8-1.2 for H/L ratio in the D-SILAC dataset, and maximum 1 replica in which the peptide was identified as substrate for the D-13MER dataset. For substrates, the enrichment threshold was 2 and above and 3 experiments in which a peptide was identified, for the D-SILAC and D-13MER sets, respectively. We note the significant difference in the number of non-substrates in the training and test sets, which preclude a rigorous comparison of performance on these sets.

### Running the calibrated protocol on the acetylome

The dataset of the human acetylome was extracted from the PhosphoSitePlus database (downloaded on 23/09/2020^34^). Hexamer peptides around sites on human proteins with at least one low-throughput experiment to support were derived with two leading and three trailing residues around the modification site. Only peptides spanning a full hexamer (*i*.*e*., the modification is not at the termini) were selected. BioGrid data^38^ for HDAC6 was downloaded on 11/09/2020 from database version 4.1.190. For pathway analysis, we used Reactome with Pathway Browser version 3.7 and database release 74. Motif prediction was done using the Eukaryotic Linear Motif resource^16^.

## Acknowledgements

This work was supported, in whole or in part, by the Israel Science Foundation, founded by the Israel Academy of Science and Humanities (grant number 717/2017 to O.S.-F.), the US-Israel Binational Science Foundation 2015207 (to O.S.-F.), and the National Institutes of Health (grant R-R01-GM-040602 to C.A.F.). J.K.V. acknowledges support from CM-SMP-KA107/426559/2019 and CM-SMP-SH/311264/2018 of the Tempus Mundi Foundation. K.D. and K.W.L acknowledge support from the Rackham Graduate School.

We thank Dr. David W. Christianson for providing us the HDAC6 construct and Dr. Nicholas J. Porter for assistance with HDAC6 purification. We are grateful to Dr. Cyril Barinka and Dr. Zsófia Kutil for sharing with us the data from their paper.

Elements of graphical abstract were drawn using figures from Servier Medical Art (https://smart.servier.com/)

## Author contributions

JKV - Investigation, Formal analysis, Writing - original draft, Visualization

KD - Investigation, Writing - original draft, Visualization

KWL - Investigation, Writing - original draft

CAF - Conceptualization, Supervision, Writing - review & editing, Funding acquisition

OFS - Conceptualization, Supervision, Writing - review & editing, Funding acquisition

## Declaration of Interests

The authors declare no competing interests.

## Notes

### Competing Interest Statement

The authors have declared no competing interest.

### Summary of Updates

Protocol calibrated with different structure and results updated accordingly, Supplementary Data added

## References

1. Inoue, A. & Fujimoto, D. Enzymatic deacetylation of histone. Biochem. Biophys. Res. Commun. 36, 146–150 (1969).

2. Allfrey, V. G., Faulkner, R. & Mirsky, A. E. Acetylation and methylation of histones and their possible role in the regulation of RNA synthesis. Proc Natl Acad Sci USA 51, 786–794 (1964).

3. Kutil, Z. et al. The unraveling of substrate specificity of histone deacetylase 6 domains using acetylome peptide microarrays and peptide libraries. FASEB J. 33, 4035–4045 (2019).

4. Hai, Y. & Christianson, D. W. Histone deacetylase 6 structure and molecular basis of catalysis and inhibition. Nat. Chem. Biol. 12, 741–747 (2016).

5. Hubbert, C. et al. HDAC6 is a microtubule-associated deacetylase. Nature 417, 455–458 (2002).

6. Yan, J. Interplay between HDAC6 and its interacting partners: essential roles in the aggresome-autophagy pathway and neurodegenerative diseases. DNA Cell Biol. 33, 567–580 (2014).

7. Bali, P. et al. Inhibition of histone deacetylase 6 acetylates and disrupts the chaperone function of heat shock protein 90: a novel basis for antileukemia activity of histone deacetylase inhibitors. J. Biol. Chem. 280, 26729–26734 (2005).

8. Kovacs, J. J. et al. HDAC6 regulates Hsp90 acetylation and chaperone-dependent activation of glucocorticoid receptor. Mol. Cell 18, 601–607 (2005).

9. Moreno-Gonzalo, O. et al. HDAC6 controls innate immune and autophagy responses to TLR-mediated signalling by the intracellular bacteria Listeria monocytogenes. PLoS Pathog. 13, e1006799 (2017).

10. Choi, S. J. et al. HDAC6 regulates cellular viral RNA sensing by deacetylation of RIG-I. EMBO J. 35, 429–442 (2016).

11. Liu, H. M. et al. Regulation of Retinoic Acid Inducible Gene-I (RIG-I) Activation by the Histone Deacetylase 6. EBioMedicine 9, 195–206 (2016).

12. Alam,x N. et al. Structure-Based Identification of HDAC8 Non-histone Substrates. Structure 24, 458–468 (2016).

13. Riester, D., Hildmann, C., Grünewald, S., Beckers, T. & Schwienhorst, A. Factors affecting the substrate specificity of histone deacetylases. Biochem. Biophys. Res. Commun. 357, 439–445 (2007).

14. Schölz, C. et al. Acetylation site specificities of lysine deacetylase inhibitors in human cells. Nat. Biotechnol. 33, 415–423 (2015).

15. Gibson, T. J., Dinkel, H., Van Roey, K. & Diella, F. Experimental detection of short regulatory motifs in eukaryotic proteins: tips for good practice as well as for bad. Cell Commun. Signal. 13, 42 (2015).

16. Kumar, M. et al. ELM-the eukaryotic linear motif resource in 2020. Nucleic Acids Res. 48, D296–D306 (2020).

17. Weatheritt, R. J., Luck, K., Petsalaki, E., Davey, N. E. & Gibson, T. J. The identification of short linear motif-mediated interfaces within the human interactome. Bioinformatics 28, 976–982 (2012).

18. Tallorin, L. et al. Discovering de novo peptide substrates for enzymes using machine learning. Nat. Commun. 9, 5253 (2018).

19. Pethe, M. A., Rubenstein, A. B. & Khare, S. D. Data-driven supervised learning of a viral protease specificity landscape from deep sequencing and molecular simulations. Proc Natl Acad Sci USA 116, 168–176 (2019).

20. Pertseva, M., Gao, B., Neumeier, D., Yermanos, A. & Reddy, S. T. Applications of machine and deep learning in adaptive immunity. Annu. Rev. Chem. Biomol. Eng. 12, 39–62 (2021).

21. Lei, Y. et al. A deep-learning framework for multi-level peptide-protein interaction prediction. Nat. Commun. 12, 5465 (2021).

22. Mahajan, S. P., Srinivasan, Y., Labonte, J. W., DeLisa, M. P. & Gray, J. J. Structural basis for peptide substrate specificities of glycosyltransferase GalNAc-T2. ACS Catal. 11, 2977–2991 (2021).

23. Chaudhury, S. & Gray, J. J. Identification of structural mechanisms of HIV-1 protease specificity using computational peptide docking: implications for drug resistance. Structure 17, 1636–1648 (2009).

24. Ollikainen, N. Flexible backbone methods for predicting and designing peptide specificity. Methods Mol. Biol. 1561, 173–187 (2017).

25. Alam, N. & Schueler-Furman, O. Modeling Peptide-Protein Structure and Binding Using Monte Carlo Sampling Approaches: Rosetta FlexPepDock and FlexPepBind. Methods Mol. Biol. 1561, 139–169 (2017).

26. London, N., Lamphear, C. L., Hougland, J. L., Fierke, C. A. & Schueler-Furman, O. Identification of a novel class of farnesylation targets by structure-based modeling of binding specificity. PLoS Comput. Biol. 7, e1002170 (2011).

27. Wolfson, N. A., Pitcairn, C. A., Sullivan, E. D., Joseph, C. G. & Fierke, C. A. An enzyme-coupled assay measuring acetate production for profiling histone deacetylase specificity. Anal. Biochem. 456, 61–69 (2014).

28. Herries, D. G. Enzyme structure and mechanism (second edition). Biochem. Educ. 13, 146 (1985).

29. Berman, H. M. et al. The protein data bank. Nucleic Acids Res. 28, 235–242 (2000).

30. Hosseinzadeh, P. et al. Anchor extension: a structure-guided approach to design cyclic peptides targeting enzyme active sites. Nat. Commun. 12, 3384 (2021).

31. Raveh, B., London, N. & Schueler-Furman, O. Sub-angstrom modeling of complexes between flexible peptides and globular proteins. Proteins 78, 2029–2040 (2010).

32. Raveh, B., London, N., Zimmerman, L. & Schueler-Furman, O. Rosetta FlexPepDock ab-initio: simultaneous folding, docking and refinement of peptides onto their receptors. PLoS ONE 6, e18934 (2011).

33. Olaoye, O. O. et al. Unique Molecular Interaction with the Histone Deacetylase 6 Catalytic Tunnel: Crystallographic and Biological Characterization of a Model Chemotype. J. Med. Chem. 64, 2691–2704 (2021).

34. Hornbeck, P. V. et al. PhosphoSitePlus, 2014: mutations, PTMs and recalibrations. Nucleic Acids Res. 43, D512–20 (2015).

35. Iaconelli, J., Xuan, L. & Karmacharya, R. HDAC6 Modulates Signaling Pathways Relevant to Synaptic Biology and Neuronal Differentiation in Human Stem-Cell-Derived Neurons. Int. J. Mol. Sci. 20, (2019).

36. Carlomagno, Y. et al. An acetylation-phosphorylation switch that regulates tau aggregation propensity and function. J. Biol. Chem. 292, 15277–15286 (2017).

37. Saito, M. et al. Acetylation of intrinsically disordered regions regulates phase separation. Nat. Chem. Biol. 15, 51–61 (2019).

38. Oughtred, R. et al. The BioGRID interaction database: 2019 update. Nucleic Acids Res. 47, D529–D541 (2019).

39. Karoutas, A. et al. The NSL complex maintains nuclear architecture stability via lamin A/C acetylation. Nat. Cell Biol. 21, 1248–1260 (2019).

40. Olety, B., Wälte, M., Honnert, U., Schillers, H. & Bähler, M. Myosin 1G (Myo1G) is a haematopoietic specific myosin that localises to the plasma membrane and regulates cell elasticity. FEBS Lett. 584, 493–499 (2010).

41. Oikonomou, K. G., Zachou, K. & Dalekos, G. N. Alpha-actinin: a multidisciplinary protein with important role in B-cell driven autoimmunity. Autoimmun. Rev. 10, 389–396 (2011).

42. Matsuyama, A. et al. In vivo destabilization of dynamic microtubules by HDAC6-mediated deacetylation. EMBO J. 21, 6820–6831 (2002).

43. Zhao, Z., Xu, H. & Gong, W. Histone deacetylase 6 (HDAC6) is an independent deacetylase for alpha-tubulin. Protein Pept. Lett. 17, 555–558 (2010).

44. Marzec, M., Eletto, D. & Argon, Y. GRP94: An HSP90-like protein specialized for protein folding and quality control in the endoplasmic reticulum. Biochim. Biophys. Acta 1823, 774–787 (2012).

45. Hart, M. J. et al. Identification of a novel guanine nucleotide exchange factor for the Rho GTPase. J. Biol. Chem. 271, 25452–25458 (1996).

46. Castañeda, C. A. et al. HDAC8 substrate selectivity is determined by long-and short-range interactions leading to enhanced reactivity for full-length histone substrates compared with peptides. J. Biol. Chem. 292, 21568–21577 (2017).

47. Vannini, A. et al. Substrate binding to histone deacetylases as shown by the crystal structure of the HDAC8-substrate complex. EMBO Rep. 8, 879–884 (2007).

48. Stiehl, D. P., Fath, D. M., Liang, D., Jiang, Y. & Sang, N. Histone deacetylase inhibitors synergize p300 autoacetylation that regulates its transactivation activity and complex formation. Cancer Res. 67, 2256–2264 (2007).

49. Black, J. C., Mosley, A., Kitada, T., Washburn, M. & Carey, M. The SIRT2 deacetylase regulates autoacetylation of p300. Mol. Cell 32, 449–455 (2008).

50. Han, Y. et al. Acetylation of histone deacetylase 6 by p300 attenuates its deacetylase activity. Biochem. Biophys. Res. Commun. 383, 88–92 (2009).

51. Hornbeck, P. V. et al. PhosphoSitePlus: a comprehensive resource for investigating the structure and function of experimentally determined post-translational modifications in man and mouse. Nucleic Acids Res. 40, D261–70 (2012).

52. Wang, F. et al. Deacetylation of FOXO3 by SIRT1 or SIRT2 leads to Skp2-mediated FOXO3 ubiquitination and degradation. Oncogene 31, 1546–1557 (2012).

53. Geng, H. et al. HIF1α protein stability is increased by acetylation at lysine 709. J. Biol. Chem. 287, 35496–35505 (2012).

54. Li, K. et al. Acetylation of WRN protein regulates its stability by inhibiting ubiquitination. PLoS ONE 5, e10341 (2010).

55. Chang, C.-W., Chuang, H.-C., Yu, C., Yao, T.-P. & Chen, H. Stimulation of GCMa transcriptional activity by cyclic AMP/protein kinase A signaling is attributed to CBP-mediated acetylation of GCMa. Mol. Cell. Biol. 25, 8401–8414 (2005).

56. Lin, R. et al. Acetylation stabilizes ATP-citrate lyase to promote lipid biosynthesis and tumor growth. Mol. Cell 51, 506–518 (2013).

57. Cohen, T. J. et al. An acetylation switch controls TDP-43 function and aggregation propensity. Nat. Commun. 6, 5845 (2015).

58. Liu, W. et al. HDAC6 regulates epidermal growth factor receptor (EGFR) endocytic trafficking and degradation in renal epithelial cells. PLoS ONE 7, e49418 (2012).

59. Wang, Z., Hu, P., Tang, F. & Xie, C. HDAC6-mediated EGFR stabilization and activation restrict cell response to sorafenib in non-small cell lung cancer cells. Med. Oncol. 33, 50 (2016).

60. Song, H. et al. Acetylation of EGF receptor contributes to tumor cell resistance to histone deacetylase inhibitors. Biochem. Biophys. Res. Commun. 404, 68–73 (2011).

61. Croft, D. et al. The Reactome pathway knowledgebase. Nucleic Acids Res. 42, D472–7 (2014).

62. Saji, S. et al. Significance of HDAC6 regulation via estrogen signaling for cell motility and prognosis in estrogen receptor-positive breast cancer. Oncogene 24, 4531–4539 (2005).

63. Li, T. et al. Histone deacetylase 6 in cancer. J. Hematol. Oncol. 11, 111 (2018).

64. Brown, N. R., Noble, M. E., Endicott, J. A. & Johnson, L. N. The structural basis for specificity of substrate and recruitment peptides for cyclin-dependent kinases. Nat. Cell Biol. 1, 438–443 (1999).

65. Daizumoto, K. et al. A DDX31/Mutant-p53/EGFR Axis Promotes Multistep Progression of Muscle-Invasive Bladder Cancer. Cancer Res. 78, 2233–2247 (2018).

66. Wee, P. & Wang, Z. Epidermal growth factor receptor cell proliferation signaling pathways. Cancers (Basel) 9, (2017).

67. Lee, Y.-S. et al. The cytoplasmic deacetylase HDAC6 is required for efficient oncogenic tumorigenesis. Cancer Res. 68, 7561–7569 (2008).

68. Qiang, L., Banks, A. S. & Accili, D. Uncoupling of acetylation from phosphorylation regulates FoxO1 function independent of its subcellular localization. J. Biol. Chem. 285, 27396–27401 (2010).

69. Ferreira, A. P. S. et al. Active glutaminase C self-assembles into a supratetrameric oligomer that can be disrupted by an allosteric inhibitor. J. Biol. Chem. 288, 28009–28020 (2013).

70. Tong, Y. et al. SUCLA2-coupled regulation of GLS succinylation and activity counteracts oxidative stress in tumor cells. Mol. Cell 81, 2303-2316.e8 (2021).

71. Dowling, C. M. et al. Multiple screening approaches reveal HDAC6 as a novel regulator of glycolytic metabolism in triple-negative breast cancer. Sci. Adv. 7, (2021).

72. New, M. et al. A regulatory circuit that involves HR23B and HDAC6 governs the biological response to HDAC inhibitors. Cell Death Differ. 20, 1306–1316 (2013).

73. Liu, J. et al. HDAC6 interacts with PTPN1 to enhance melanoma cells progression. Biochem. Biophys. Res. Commun. 495, 2630–2636 (2018).

74. Hsu, F. F. et al. Acetylation is essential for nuclear heme oxygenase-1-enhanced tumor growth and invasiveness. Oncogene 36, 6805–6814 (2017).

75. Zhu, Y. et al. Dynamic regulation of ME1 phosphorylation and acetylation affects lipid metabolism and colorectal tumorigenesis. Mol. Cell 77, 138-149.e5 (2020).

76. Skultetyova, L. et al. Human histone deacetylase 6 shows strong preference for tubulin dimers over assembled microtubules. Sci. Rep. 7, 11547 (2017).

77. Katz, C. et al. Studying protein-protein interactions using peptide arrays. Chem. Soc. Rev. 40, 2131–2145 (2011).

78. Kim, S.-Y. et al. An Alternative Strategy for Pan-acetyl-lysine Antibody Generation. PLoS ONE 11, e0162528 (2016).

79. Virág, D. et al. Current Trends in the Analysis of Post-translational Modifications. Chromatographia 83, 1–10 (2020).

80. Tropea, J. E., Cherry, S. & Waugh, D. S. Expression and purification of soluble His(6)-tagged TEV protease. Methods Mol. Biol. 498, 297–307 (2009).

81. Huang, X. & Hernick, M. A fluorescence-based assay for measuring N-acetyl-1-D-myo-inosityl-2-amino-2-deoxy-α-D-glucopyranoside deacetylase activity. Anal. Biochem. 414, 278–281 (2011).

82. Alford, R. F. et al. The Rosetta All-Atom Energy Function for Macromolecular Modeling and Design. J. Chem. Theory Comput. 13, 3031–3048 (2017).

83. Krystkowiak, I., Manguy, J. & Davey, N. E. PSSMSearch: a server for modeling, visualization, proteome-wide discovery and annotation of protein motif specificity determinants. Nucleic Acids Res. 46, W235–W241 (2018).

84. Vacic, V., Iakoucheva, L. M. & Radivojac, P. Two Sample Logo: a graphical representation of the differences between two sets of sequence alignments. Bioinformatics 22, 1536–1537 (2006).

85. Robin, X. et al. pROC: an open-source package for R and S+ to analyze and compare ROC curves. BMC Bioinformatics 12, 77 (2011).

