## Supplementary for "Structure-based prediction of HDAC6 substrates validated by enzymatic assay reveals determinants of promiscuity and detects new potential substrates"

for

#### Contents

**Supplementary Figure 1. Performance of selected protocol on different starting structures.**	**2**

**Supplementary Figure 2. Position specific scoring matrices of the datasets.**	**3**

**Supplementary Figure 3. PSSM scores and experimental values do not correlate.**	**4**

**Supplementary Figure 4. Position-wise correlation of PSSMs derived from high-throughput experiments.**	**5**

**Supplementary Table 1. Non-hexameric and inaccurately measured peptides.**	**8**

**Supplementary Table 2. Datasets evaluated in this study.**	**9**

**Supplementary Table 3. Kinetic parameters of H3 K14Ac peptide and full-length protein.**	**10**

**Supplementary Table 4. Performance of protocol on high-throughput datasets.**	**11**

**Supplementary Table 5. Files of Rosetta protocols.**	**12**

**Supplementary Table 6. Commands and flags of Rosetta runs.**	**13**

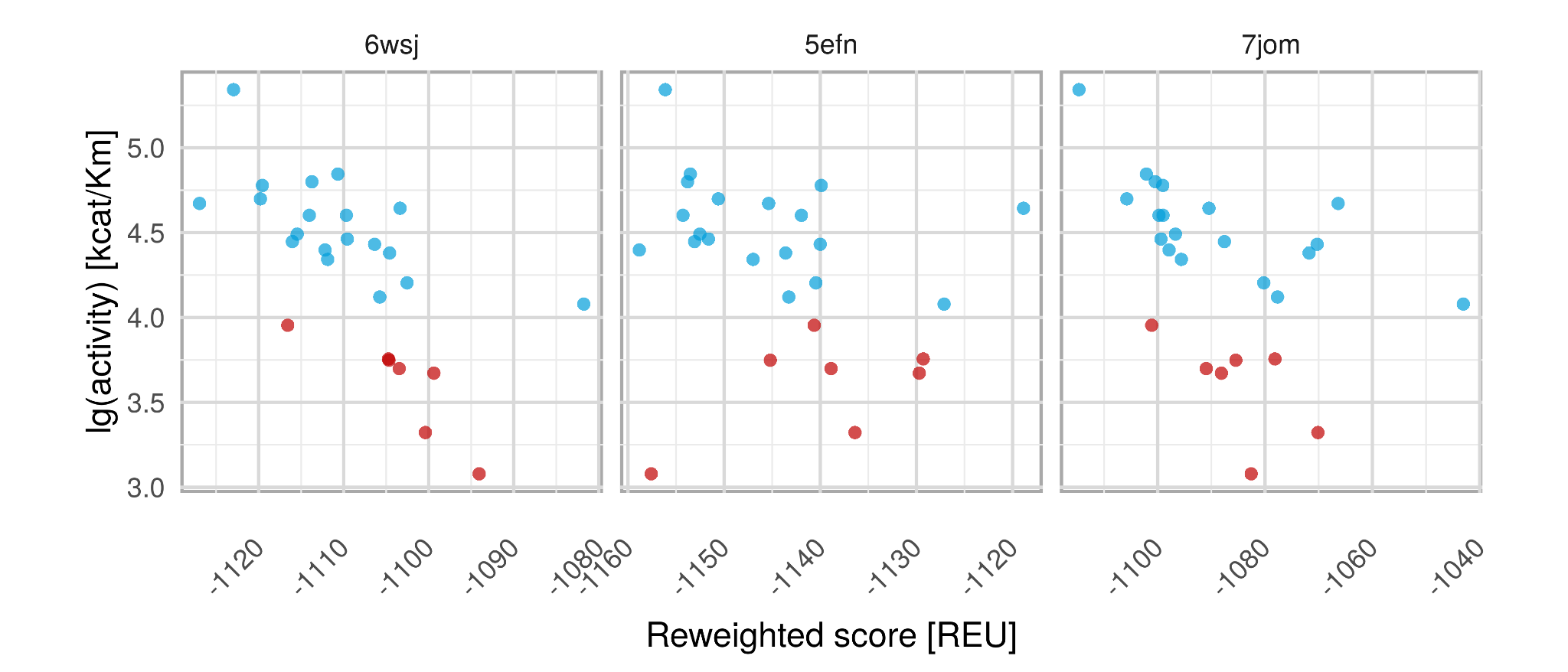

###### **Supplementary Figure S1****. Performance on different starting structures**

Shown are results for the training set (D-TRAINING). The calibrated protocol was run on a structure of HDAC6 DD2 domain bound to (1) a cyclic peptide (6WSJ; same as **Figure 2A**, (2) a trimer peptide connected to coumarine (5EFN), and (3) a small molecule inhibitor (7JOM). (blue: substrates; red: non-substrates).

| A  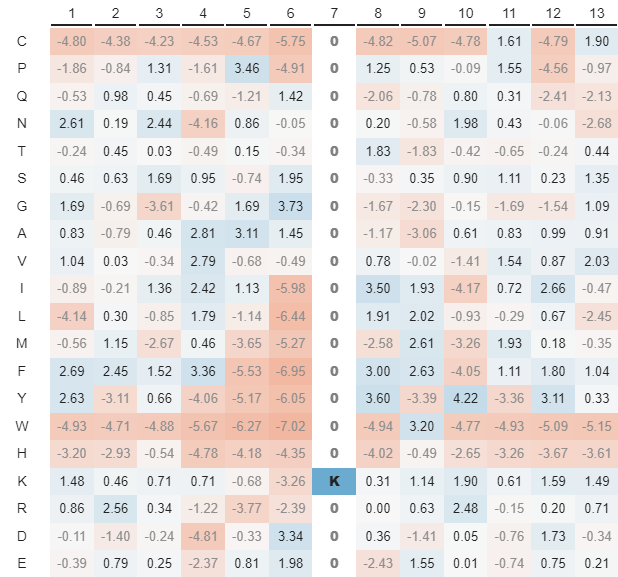 | C  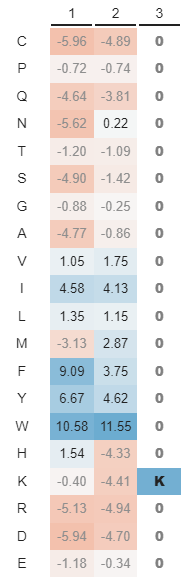 |
| --- | --- |
| B  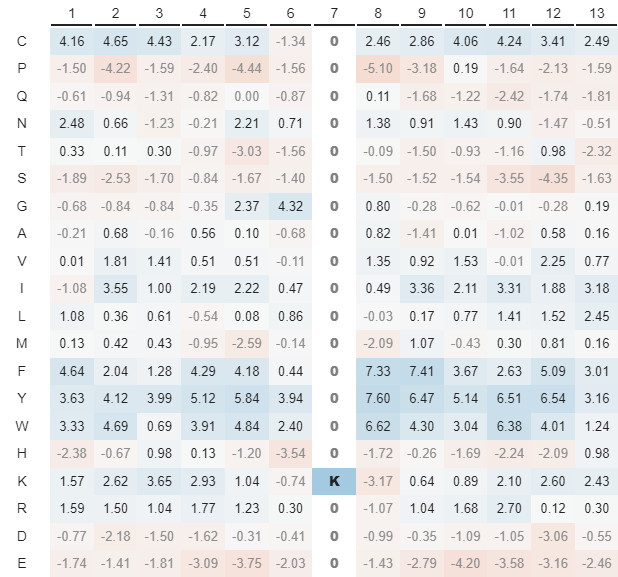 | D  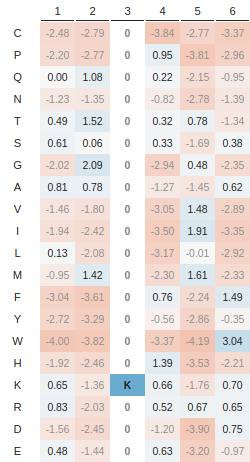 |
| **Supplementary Figure S2****. Position specific scoring matrices derived from different datasets**  PSSMS were generated from **A)** D-SILAC, **B)** D-13MER, **C)** D-3MER, and **D)** D-TRAINING sets by applying PSSMSearch to the substrate list of each experiment (for definition of substrates, see **Supplementary Table S1**). (blue: enriched residues, red: depleted amino acids). | |

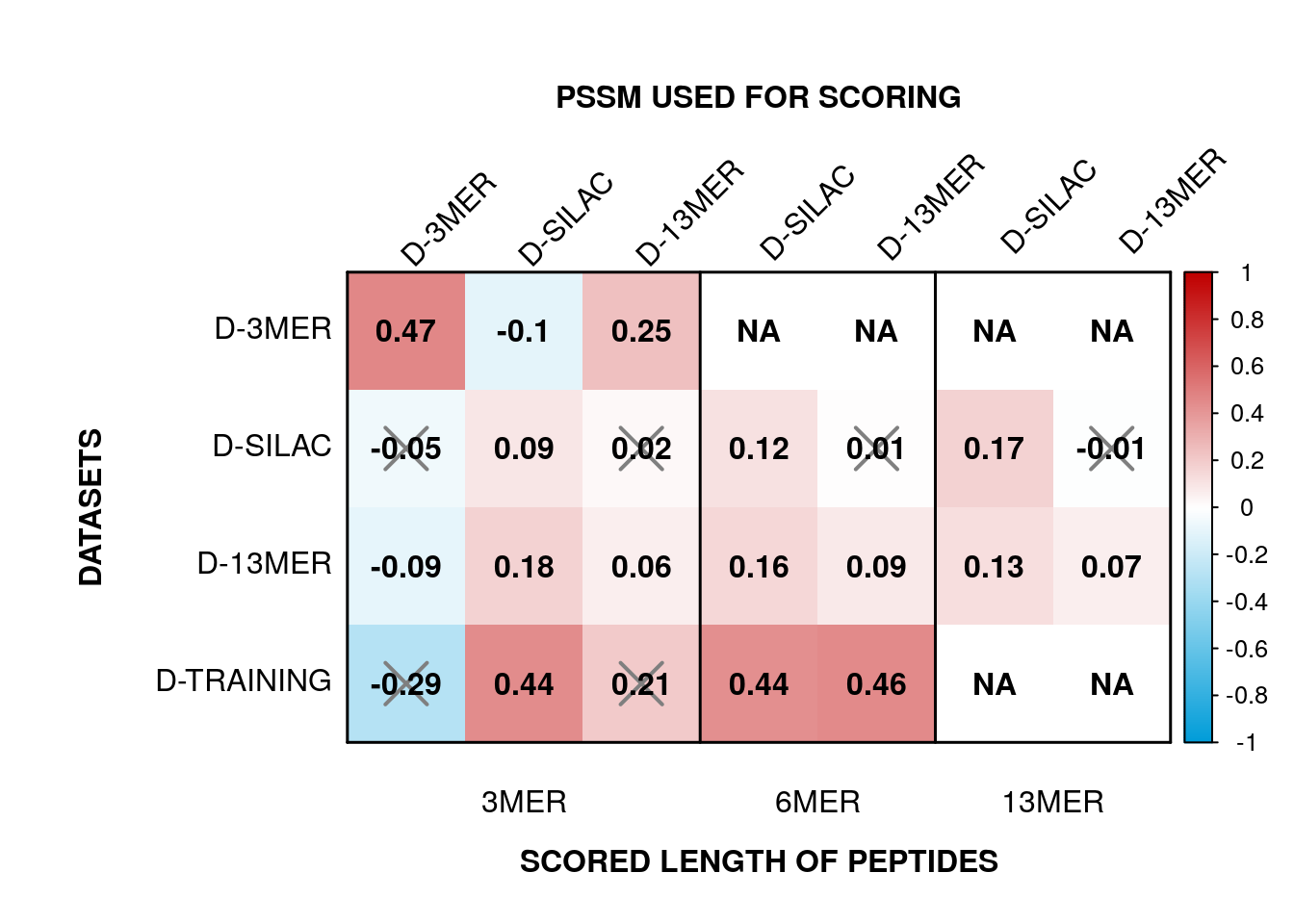

###### **Supplementary Figure S3****. PSSM scores and experimental values do not correlate.**

Each dataset was cross-scored with every obtained PSSM depicted in **Supplementary Figure S2**. Three different lengths for scoring (3-, 6- and 13-mers) were used where applicable. D-3MER, D-SILAC and D-13MER labels at the top represent the dataset that the respective PSSM was derived from. 3MER, 6MER and 13MER labels indicate the number of amino acids used for scoring. X indicates non-significant correlations.

A
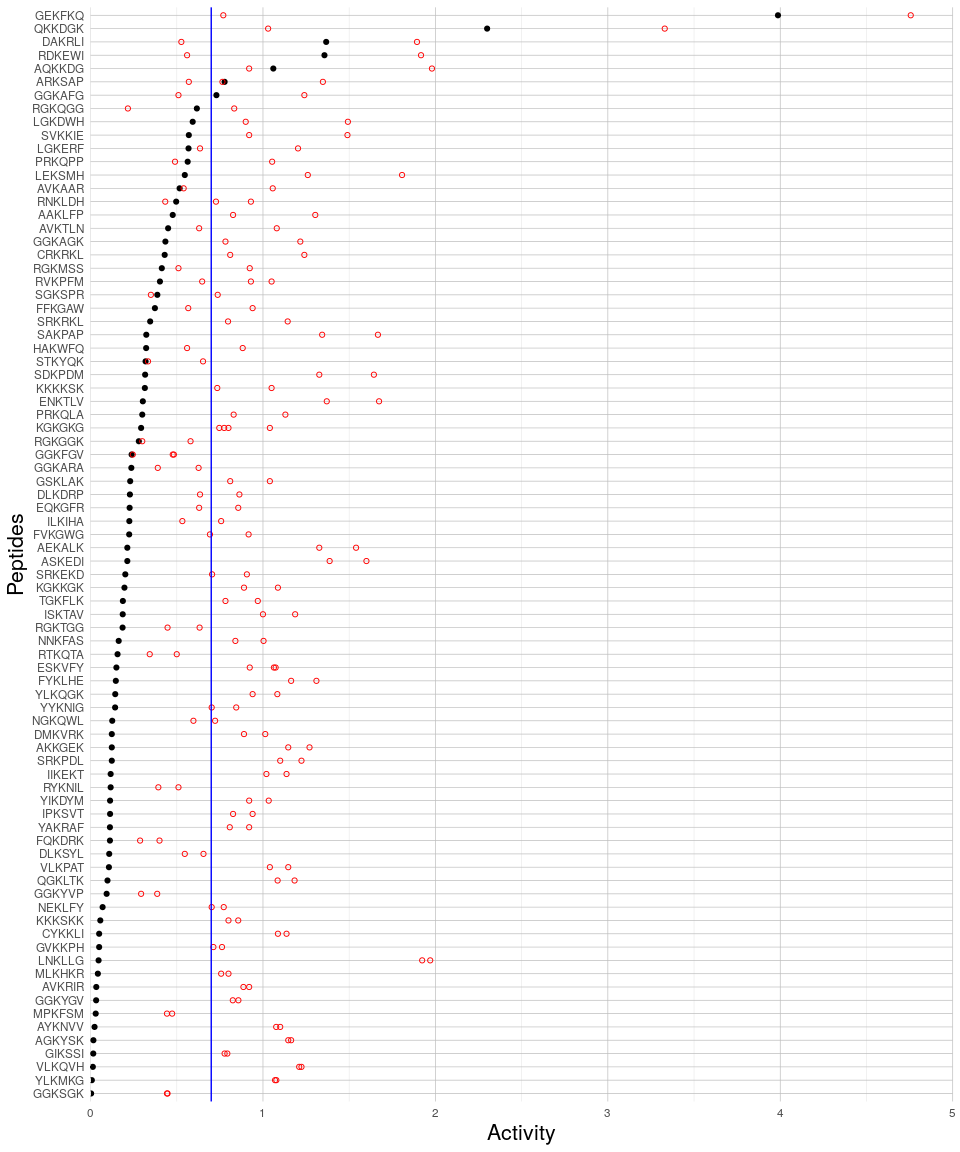

B

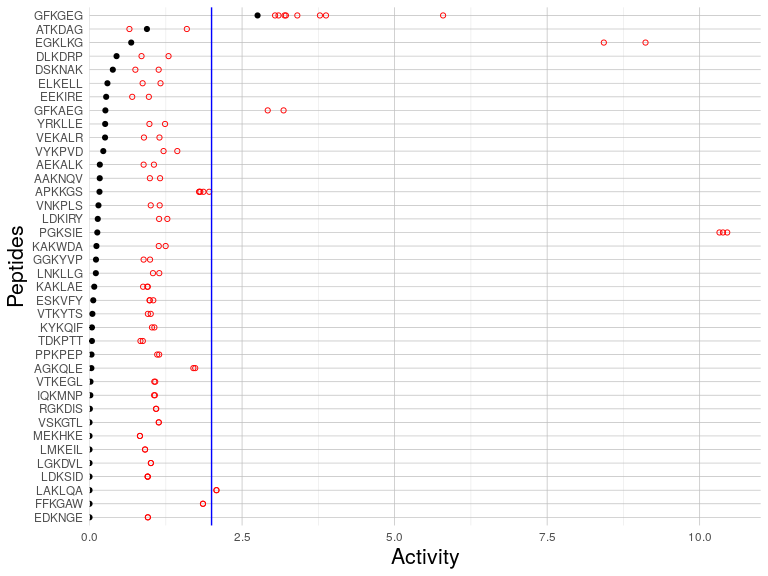

###### **Supplementary Figure S4****. Range of measured activities for peptides sharing identical core hexamers with differing flanking regions.**

**A)** D-13MER and **B)** D-SILAC datasets. (Black dots: range, red circles: measured values, blue line: threshold of substrate-non-substrate distinction) Related to [**Figure 4**](https://docs.google.com/document/d/1VQqFC-0O9emiuj7rcuS3ttDjD2CvG_CZM2OUVi4fYyM#fig_comparison).

###### **Supplementary Table S1****.** **Datasets evaluated in this study.** The unit of measurement of substrate activity and parameters used to define substrates are indicated. See Text for more details. Related to [**Figure 4**](https://docs.google.com/document/d/1VQqFC-0O9emiuj7rcuS3ttDjD2CvG_CZM2OUVi4fYyM#fig_comparison).

| **Dataset [reference]** | **N***^b^****_peptides_*** | **N*_substrates_*** | **N*_non-substrates_*** | **Cutoff for substrates** | **Cutoff for non-substrates** | **Unit** |
| --- | --- | --- | --- | --- | --- | --- |
| D-TRAINING*^a^* | 26 | 19 | 7 | ≧10^4^ | - | *k*_cat_/*K*_M_ |
| D-CAPPED*^a^* | 16 | 16 | 0 | ≧10^4^ | - | *k*_cat_/*K*_M_ |
| D-13MER[^1^](https://sciwheel.com/work/citation?ids=8374723&pre=&suf=&sa=0) | 6797 | 395 (51)*^c^* | 4127 (245)*^c^* | 4 out of 4 | 1 out of 4 | # experiments that identified as substrate |
| D-GCMS[^1^](https://sciwheel.com/work/citation?ids=8374723&pre=&suf=&sa=0) | 24 | 17 | 7 | - | - | - |
| D-SILAC [^2^](https://sciwheel.com/work/citation?ids=1577041&pre=&suf=&sa=0) | 929 | 63 (18)*^c^* | 328 (278)*^c^* | ≧2 | ≦1 | ratio (H/L) |
| D-3MER [^3^](https://sciwheel.com/work/citation?ids=10264624&pre=&suf=&sa=0) | 361 | 35 | 33 | ≧70 | ≦21 | - (intensity) |
| *^a^* D-TRAINING & D-CAPPED: present study (see [**Table 1**](https://docs.google.com/document/d/1VQqFC-0O9emiuj7rcuS3ttDjD2CvG_CZM2OUVi4fYyM#tab_exp)**)**.  *^b^ N* indicates the number of respective peptides  *^c^* In parentheses: peptides that were reported both in the D-13MER and D-SILAC sets. | | | | | | |

###### **Supplementary Table S2****. Kinetic parameters of the H3 K14_Ac_ peptide and full-length protein from which it was derived.** Related to **Table 1.**

| **H3 K14ac substrate** | ***k*_cat_/*K*_M_**  **(M^-1^s^-1^)** | ***k*_cat_**  **(s^-1^)** | ***K*_M_**  **(μM)** |
| --- | --- | --- | --- |
| 13-mer: RKSTGG(K-ac)APRKQL | 140,000$\pm$10,000 | 4.1 ± 0.3 | 28 ± 5 |
| Full length protein | 80,000$\pm$40,000 | 1.4 ± 0.6 | 20 ± 10 |

###### **Supplementary Table S3****.** **Peptides measured for HDAC6 deacetylation in this study but not used for training or validation.**

These peptides were not included due to poor measurement accuracy, or peptide length beyond 6 residues. *calculated from only 3 data points. Related to [**Table 1**](https://docs.google.com/document/d/1VQqFC-0O9emiuj7rcuS3ttDjD2CvG_CZM2OUVi4fYyM#tab_exp).

| **Peptide** | **Protein (site of modification)** | ***k*_cat_/*K*_M_**  **(M^-1^s^-1^)** | ***k*_cat_**  **(s^-1^)** | ***K*_M_**  **(μM)** | **Rosetta reweighted score [REU]** |
| --- | --- | --- | --- | --- | --- |
| ME(K-Ac)KKE | GBP7 (K-389) | 3,300 ± 200* | >0.3* | >150* | -1,105 |
| QD(K-Ac)PLR | CCDC86 (K-261) | >2,000* | 0.11 ± 0.04* | <50* | -1,092 |
| kgGA(K-Ac)RHR | H4K16 (K-16)[^4^](https://sciwheel.com/work/citation?ids=10272230&pre=&suf=&sa=0) | 70,000 ± 20,000 | 1.23 ± 0.09 | 17 ± 6 | -1,106 |
| SLG(K-Ac)DWHR | CRIP1 (K-22), CRIP1 (K-144) | 31,000 ± 5,000 | 10 ± 2 | 300 ± 200 | -1,113 |
| rMF(K-Ac)QFNK | TRIM28 (K-770) | 21,000 ± 1,000 | >2 | >200 | -1,110 |
| riIL(K-Ac)ASR | MSH2 (K-635)[^5^](https://sciwheel.com/work/citation?ids=10264623&pre=&suf=&sa=0) | 20,000 ± 3,000 | 4 ± 1 | 180 ± 80 | -1,115 |

###### **Supplementary Table S4****. Performance of protocol on high-throughput datasets.**

(MCC: Matthews correlation coefficient, AUC: area under the ROC curve)

| **Dataset** | **Specificity** | **Sensitivity** | **MCC** | **Spearman correlation** | **AUC** |
| --- | --- | --- | --- | --- | --- |
| **D-13MER** | 0.18 | 0.77 | -0.05 | -0.01 | 0.49 |
| **D-SILAC** | 0.19 | 0.89 | 0.05 | -0.05 | 0.52 |

###### **Supplementary Table S5****. Files of Rosetta protocols.**

Constraint File for docking and minimization measured on PDB ID 6WSJ. (Chain A: receptor, chain F is the peptide). Fixbb resfile is used for peptide threading.

| **Constraint file** | Dihedral N 3F CA 3F C 3F N 4F CIRCULARHARMONIC -36.7 0.5  AtomPair OG 531A N 3F HARMONIC 2.9 0.2  AtomPair OD2 705A OH 3F HARMONIC 3.7 0.2  AtomPair OD1 705A OH 3F HARMONIC 5.2 0.2  AtomPair OD2 612A OH 3F HARMONIC 3.4 0.2  AtomPair OD1 612A OH 3F HARMONIC 3.7 0.2  AtomPair ND1 614A OH 3F HARMONIC 3.8 0.2  AtomPair O 582A NZ 3F HARMONIC 3.3 0.2  AtomPair CE1 583A CD 3F HARMONIC 4.0 0.2  AtomPair CE2 583A CD 3F HARMONIC 3.7 0.2  AtomPair CD2 643A CG 3F HARMONIC 4.0 0.2  AtomPair CD1 643A CG 3F HARMONIC 3.7 0.2 |
| --- | --- |
| **fixxb resfile** | NATRO  start  1 F PIKAA E EX 1 EX 2 USE_INPUT_SC  2 F PIKAA G EX 1 EX 2 USE_INPUT_SC  4 F PIKAA F EX 1 EX 2 USE_INPUT_SC  5 F PIKAA V EX 1 EX 2 USE_INPUT_SC  6 F PIKAA R EX 1 EX 2 USE_INPUT_SC |

###### **Supplementary Table S6****. Commands and flags of Rosetta runs.**

All commands were run with Rosetta v2020.28.

| **Prepack** | $ROSETTA_HOME/main/source/bin/FlexPepDocking.default.linuxgccrelease -s input/6wsj.pdb -ex1 -ex2aro -use_input_sc -flexpep_prepack -nstruct 1 -scorefile ppk.score.sc -flexpep_score_only -out:path:pdb input -out:path:score output -unboundrot input/5eem.pdb 6wsj.pdb |
| --- | --- |
| **Peptide docking** | $ROSETTA_HOME/main/source/bin/FlexPepDocking.mpi.linuxgccrelease -s input/6wsj.ppk.pdb -ex1 -ex2aro -use_input_sc -constraints:cst_fa_file input/constraints_6wsj.cst -constraints:cst_fa_weight 1.0 -unboundrot input/5eem.pdb input/6wsj.ppk.pdb -nstruct 250 -flexpep_score_only -scorefile refine.score.sc -out:path:pdb output -out:file:silent output/decoys.silent -out:file:silent_struct_type binary -out:path:score output -overwrite -flexPepDocking:pep_refine -lowres_preoptimize  (-min_receptor_bb flag was added in some protocols, as described in Methods) |
| **Peptide threading** | $ROSETTA_HOME/main/source/bin/fixbb.default.linuxgccrelease -database $ROSETTA_HOME/database -resfile resfile -s template.pdb -ex1 -ex2aro -use_input_sc -scorefile design.score.sc -nstruct 1 -unboundrot 5eem.pdb template.pdb |
| **Minimization** | $ROSETTA_HOME/main/source/bin/FlexPepDocking.mpi.linuxgccrelease -s start.ppk.pdb -ex1 -ex2 -ex3 -ex4 -constraints:cst_fa_file constraints.cst -constraints:cst_fa_weight 1.0 -scorefile min.score.sc -unboundrot 5eem.pdb start.ppk.pdb -flexPepDockingMinimizeOnly -flexpep_score_only  (-min_receptor_bb flag was added in some protocols, as described in Methods) |

### References

[1.	Kutil, Z. *et al.* The unraveling of substrate specificity of histone deacetylase 6 domains using acetylome peptide microarrays and peptide libraries. *FASEB J.* **33**, 4035–4045 (2019).](https://sciwheel.com/work/bibliography/8374723)

[2.	Schölz, C. *et al.* Acetylation site specificities of lysine deacetylase inhibitors in human cells. *Nat. Biotechnol.* **33**, 415–423 (2015).](https://sciwheel.com/work/bibliography/1577041)

[3.	Riester, D., Hildmann, C., Grünewald, S., Beckers, T. & Schwienhorst, A. Factors affecting the substrate specificity of histone deacetylases. *Biochem. Biophys. Res. Commun.* **357**, 439–445 (2007).](https://sciwheel.com/work/bibliography/10264624)

[4.	Sui, L., Huang, R., Yu, H., Zhang, S. & Li, Z. Inhibition of HDAC6 by tubastatin A disrupts mouse oocyte meiosis via regulating histone modifications and mRNA expression. *J. Cell. Physiol.* **235**, 7030–7042 (2020).](https://sciwheel.com/work/bibliography/10272230)

[5.	Zhang, M. *et al.* HDAC6 deacetylates and ubiquitinates MSH2 to maintain proper levels of MutSα. *Mol. Cell* **55**, 31–46 (2014).](https://sciwheel.com/work/bibliography/10264623)
